# Assessing connectivity despite high diversity in island populations of a malaria mosquito

**DOI:** 10.1101/430702

**Authors:** Christina M. Bergey, Martin Lukindu, Rachel M. Wiltshire, Michael C. Fontaine, Jonathan K. Kayondo, Nora J. Besansky

## Abstract

Documenting isolation is notoriously difficult for species with vast polymorphic populations. High proportions of shared variation impede estimation of connectivity, even despite leveraging information from many genetic markers. We overcome these impediments by combining classical analysis of neutral variation with assays of the structure of selected variation, demonstrated using populations of the principal African malaria vector *Anopheles gambiae*. Accurate estimation of mosquito migration is crucial for efforts to combat malaria. Modeling and cage experiments suggest that mosquito gene drive systems will enable malaria eradication, but establishing safety and efficacy requires identification of isolated populations in which to conduct field-testing. We assess Lake Victoria islands as candidate sites, finding one island 30 kilometers offshore is as differentiated from mainland samples as populations from across the continent. Collectively, our results suggest sufficient contemporary isolation of these islands to warrant consideration as field-testing locations and illustrate shared adaptive variation as a useful proxy for connectivity in highly polymorphic species.

---

The difficulties in estimating migration with genetic methods are exacerbated for large, interconnected populations exhibiting shallow population structure. Large population sizes result in high levels of polymorphism in the genome and impede accurate estimation of connectivity [1] and discernment of demographic independence from panmixia [2]. Population genetic methods for estimating migration using neutral markers may thus have limited utility when such a high proportion of diversity is shared between populations, a failing that is only partially redressed with the high quantity of markers available from massively parallel sequencing. The most powerful window into migration may instead be the distribution of selected variants [3].

The major African malaria vector *Anopheles gambiae sensu stricto* (henceforth *An. gambiae*) is among the most genetically diverse eukaryotic species [4], with shallow population structure [4, 5] that complicates efforts to estimate connectivity from genetic data. Overcoming these obstacles to infer migration accurately is crucial for control efforts to reduce the approximately 445,000 annual deaths attributable to malaria [6]. Such vector control efforts include novel methods involving the release of genetically modified mosquitoes. The most promising involve introducing transgenes into the mosquito genome or its endosymbionts that interrupt pathogen transmission coupled with a gene drive system to propagate the effector genes through a population [7–9]. Such systems have recently been successfully engineered in the laboratory [10]. A detailed understanding of population structure and connectivity is essential for effective implementation of any genetic control method, not least a gene drive system designed to spread in a super-Mendelian fashion.

Here, we analyze population structure, demographic history, and migration between populations from genome-wide variation in *An. gambiae* mosquitoes living near and on the Ssese archipelago of Lake Victoria in Uganda (Fig. 1). We augment these analyses with a demonstration of our framework using selective sweep sharing as a proxy for connectivity when inferring migration in taxa with high variation. Islands present natural laboratories for disentangling the determinants of population structure, as gene flow—likely important in post-dry season recolonization [11]—is reduced. In addition to the high malaria prevalence of the islands (44% in children; 30% in children country-wide; [12]), we were motivated by the potential of such an island to be a field site for future tests of gene-drive vector control strategies: Geographically-isolated islands have been proposed as locales to test the dynamics of transgene spread while limiting their movement beyond the study population [13–16]. Antecedent studies of population structure and connectivity of potential release sites are crucial to evaluate the success of such field trials, as well as to quantify the chance of migration of transgenic insects carrying constructs designed to propagate across mosquito populations and country borders.

**Figure 1:**
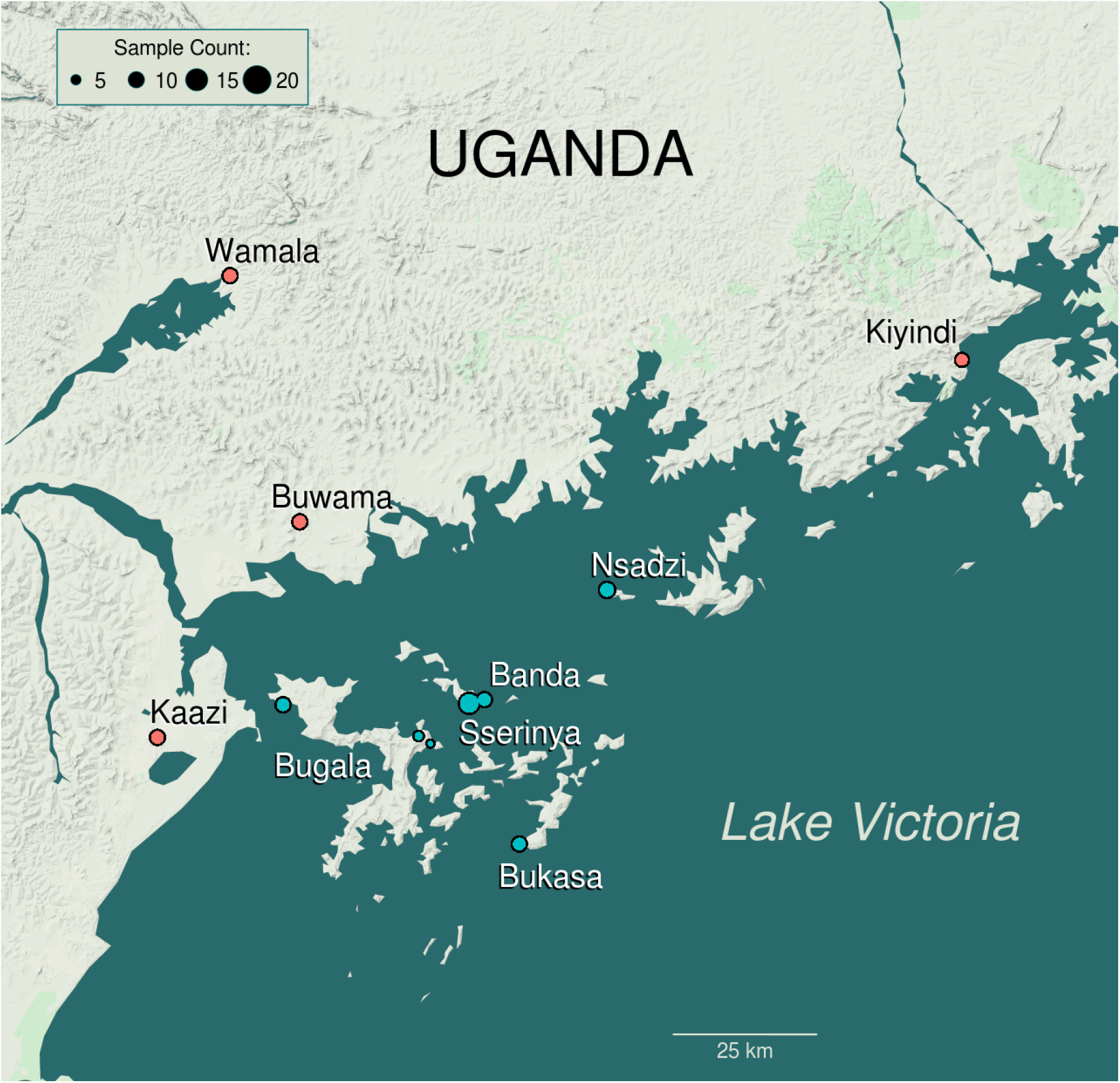
Map of Lake Victoria Basin study area. Map of study area showing sampling localities on Ssese Islands (blue) and mainland localities (red) in Lake Victoria Basin. The Ag1000G reference population, Nagongera, Tororo District, is not shown, but lies 111 km NE of Kiyindi, 57 km from the shore of Lake Victoria. Map data copyright 2018 Google.

## Results

The Ssese Islands are approximately 4-50 km from the mainland, and vary in size, infrastructure, and accessibility. Sampled islands range from Banda—a small, largely forested island of approximately 1 square kilometer with a single settlement—to Bugala—296 square kilometers, site of a 10,000 ha oil palm plantation [17], and linked to the mainland via ferry service [18]. To explore the partitioning of *An. gambiae* genetic variation in the Lake Victoria Basin (LVB), we sequenced the genomes of 116 mosquitoes from 5 island and 4 mainland localities (Fig. 1, Supplementary Table S1). We sequenced 10-23 individuals per site to an average depth of 17.6 *±* 4.6 (Supplementary Table S2). After filtering (detailed in Methods), we identified 28.6 million high quality Single Nucleotide Polymorphisms (SNPs). We merged our dataset with that of the *An. gambiae* 1000 Genomes project (Ag1000G; [4]) for a combined dataset of 12.54 million SNPs (9.86 million after linkage disequilibrium pruning) in 881 individuals.

## Genetic structure

We analyzed LVB population structure with context from continent-wide populations [4] of *An. gambiae* and sister species *Anopheles coluzzii* mosquitoes (formerly known as *An. gambiae* M molecular form [19]). Both Bayesian clustering ([20]; Fig. 2a) and principal component analysis (PCA; Supplementary Fig. S1) showed LVB individuals closely related to the Ugandan reference population (Nagongera, Tororo; 0*°*46*′*12.0*″*N, 34*°*01*′*34.0*″*E; *∼*57 km from Lake Victoria; Fig. 1). With *≥* 6 clusters (which optimized predictive accuracy in the clustering analysis; Supplementary Fig. S2), island samples had distinct ancestry proportions (Fig. 2a), and with *k* = 9 clusters, we observed additional subdivision in LVB samples and the assignment of the majority of Ssese individuals’ ancestry to a largely island-specific component (Figs. 2a, 2b, and Supplementary Fig. S3).

**Figure 2:**
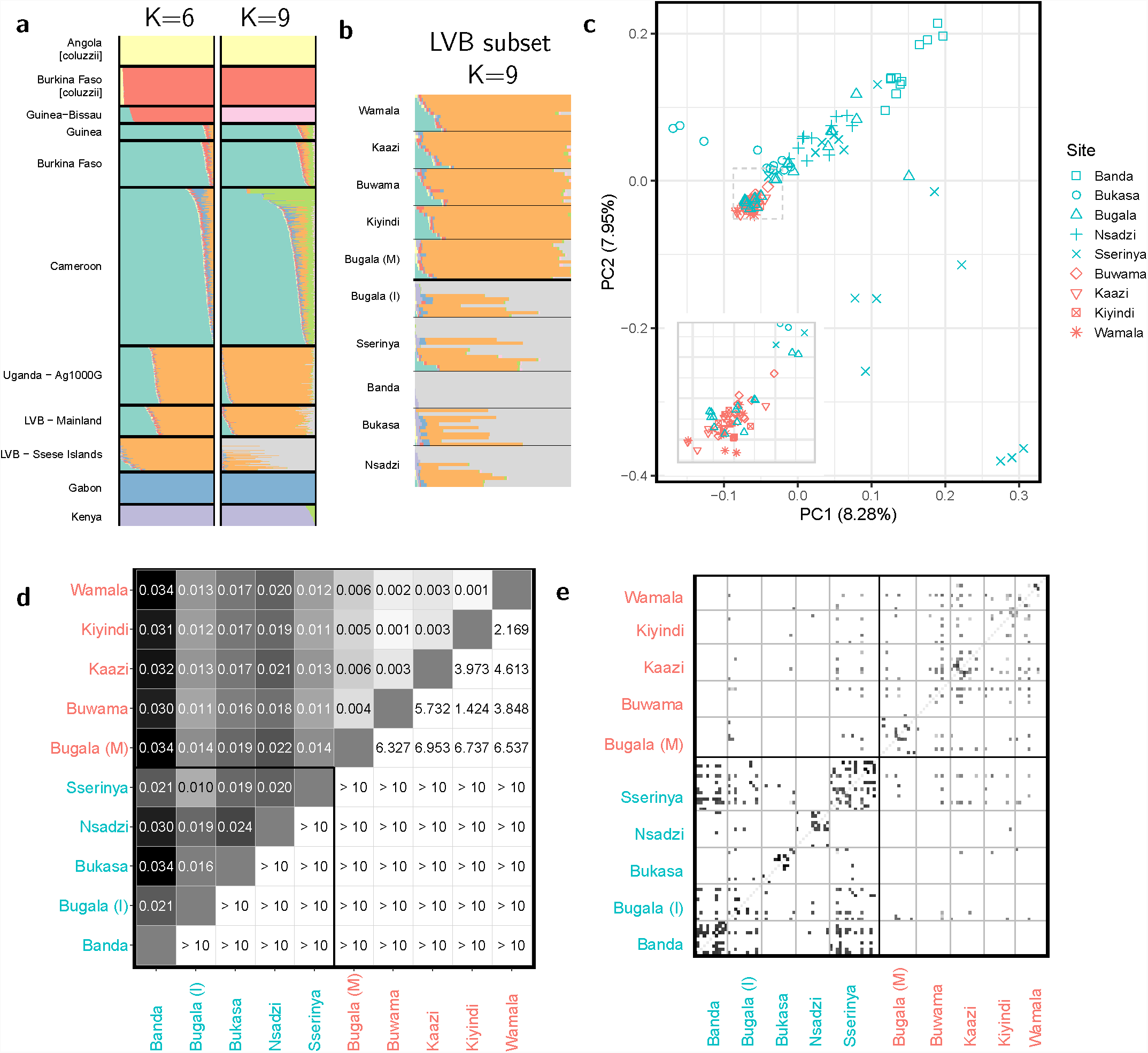
Population structure in the Lake Victoria Basin. Analyses are based on chromosome 3 to avoid segregating inversions on other chromosome, unless otherwise noted. (A) ADMIXTURE-inferred ancestry of individuals in Lake Victoria Basin. Results based on analysis of LVB and Ag1000G merged dataset. Analysis is restricted to *A. gambiae s. s.*. Clustering shown for *k* = 6 clusters, which minimizes cross validation error, and *k* = 9 clusters, the lowest *k* for which island individuals have the majority of their ancestry assigned to an island-specific cluster. (B) Results of the clustering analysis with *k* = 9 clusters for LVB individuals, split by sampling locality. (C) Plot of first two components of PCA of Lake Victoria Basin individuals showing locality of origin. Mainland individuals are colored red, while island individuals are blue, and point shape indicates sampling locality. Based on these results and that of ADMIXTURE analysis, the island sample of Bugala was split into mainlandand island-like subpopulations (“Bugala (M)” and “Bugala (I),” respectively) for subsequent analyses (Fig. S4). (D) Heatmap of *F*_*ST*_ between sites (lower triangle) and associated *z*-score computed via block jackknife (upper triangle). “Bugala (M)” and “Bugala (I)” are the mainlandand island-like subpopulations of Bugala. (E) Proportion of genome-wide pairwise IBD sharing between individuals, based on the full genome. Each small square represents a comparison between two individuals, and darker colors indicate a higher proportion of the two genomes is in IBD, shaded on a logarithmic scale. Individuals are grouped by locality.

PCA of only LVB individuals indicated little differentiation among mainland samples in the first two components and varying degrees of differentiation on islands, with Banda, Sserinya, and Bukasa the most extreme (Fig. 2c). Twelve of 23 individuals from Bugala, the largest, most developed, and most connected island, exhibited affinity to mainland individuals instead of ancestry typical of the islands (Supplementary Fig. S4). As both PCA and clustering analyses revealed this differentiation, we split the Bugala sample into mainland- and island-like subsets for subsequent analyses (hereafter referenced as “Bugala (M)” and “Bugala (I),” respectively). Individuals with partial ancestry attributable to the component prevalent on the mainland and the rest to the island-specific component were present on all islands except Banda.

Differentiation concurred with observed population structure. Mean *F*_*ST*_ between sampling localities (range: 0.001-0.034) was approximately 0 (*≤* 0.003) for mainland-mainland comparisons and was highest in comparisons involving the small island Banda (Fig. 2d). Geographic distances and *F*_*ST*_ were uncorrelated (Mantel *p* = 0.88; Supplementary Fig. S5). Island samples showed greater within- and between-locality sharing of genomic regions identical by descent (IBD), with sharing between nearby islands Sserinya, Banda and Bugala (Fig. 2e). Importantly, Banda Island shared no IBD regions with mainland sites, underscoring its contemporary isolation from the mainland.

## Genetic diversity

Consistent with the predicted decrease in genetic variation for semi-isolated island populations due to inbreeding and smaller effective population sizes (*N*_*e*_), islands displayed slightly lower nucleotide diversity (*π*; Wilcoxon rank sum test *p <* 0.001; Fig. 3a), a higher proportion of shared to rare variants (Tajima’s *D*; *p <* 0.001; Fig. 3b), and more linkage among SNPs (LD; *r*^2^; *p <* 0.001; Fig. 3f). They were however, similar in inbreeding coefficient (*F*; *p* = 0.0719; Fig. 3c), number of long runs of homozygosity (*F*_*ROH*_; *p* = 0.182; Fig. 3d), and proportions of low frequency SNPs (Fig. 3e). The small island Banda was the most extreme in these measures.

**Figure 3:**
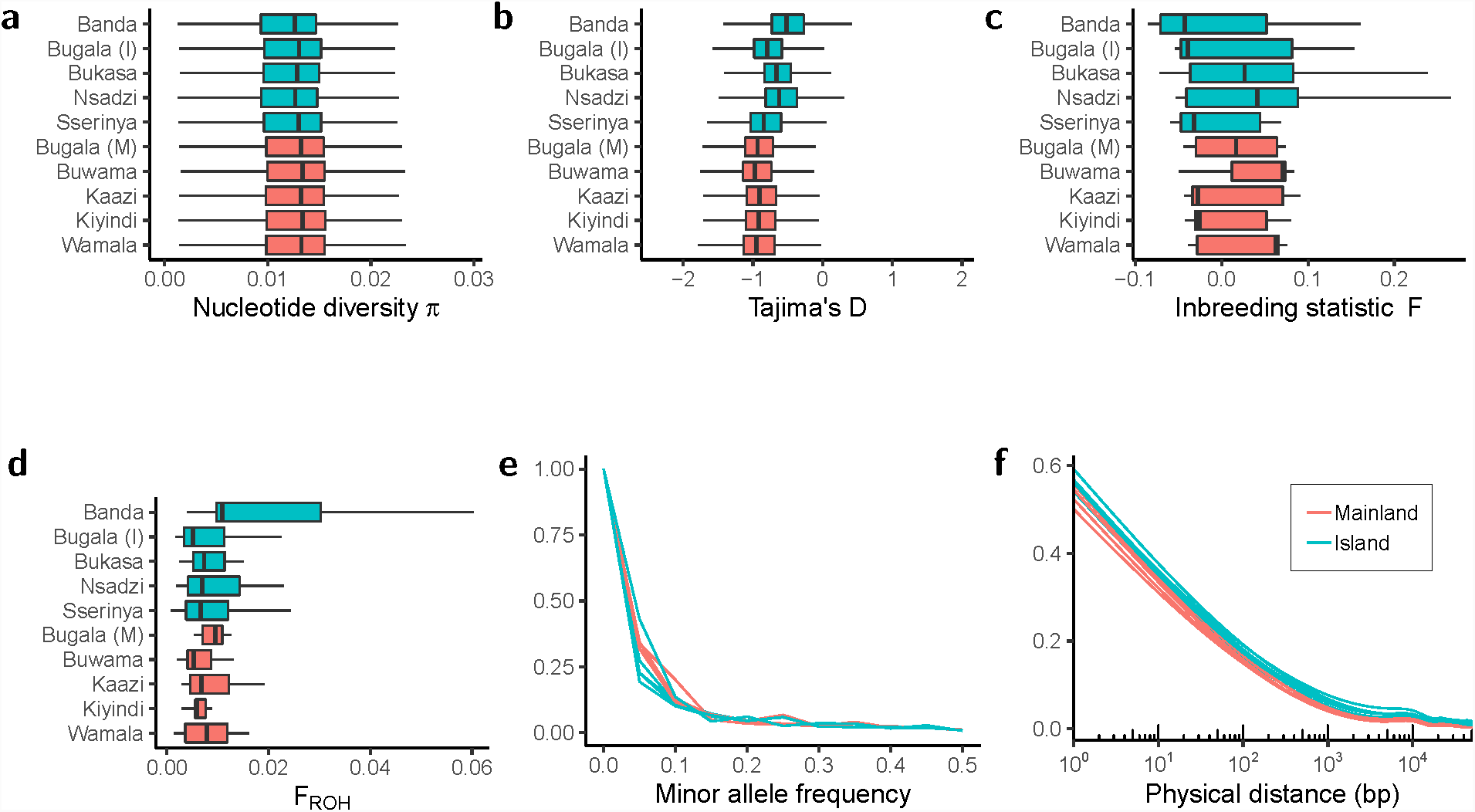
Diversity metrics in the Lake Victoria Basin samples. Shown are a (A) boxplot of nucleotide diversity (*π*; in 10 kilobase windows), (B) boxplot of Tajima’s *D* (in 10 kilobase windows), (C) boxplot of inbreeding statistic (*F*), (D) boxplot of length of runs of homozygosity (*F*_*ROH*_), (E) histogram of Minor Allele Frequency (MAF), and (F) decay in linkage disequilibrium (*r*^2^), all grouped by sampling locality. For all boxplots, outlier points are not shown.

## Demographic history

To test islands for isolation and demographic independence from the mainland, we inferred the population history of LVB samples by estimating long-term and recent trends in *N*_*e*_ using stairway plots [21] based on the site frequency spectrum (SFS; Fig. 4a) and patterns of IBD sharing ([22]; Fig. 4b), respectively. Short-term final mainland sizes were unrealistically high, likely due to low samples sizes for each locality, but island-mainland differences were nonetheless informative. In both, islands had consistently lower *N*_*e*_ compared to mainland populations extending back 500 generations (*∼* 50 years) and often severely fluctuated, particularly in the last 250 generations (*∼* 22 years). Mainland sites Wamala and Kaazi had island-like recent histories, with Wamala abruptly switching to an island-like pattern around 2005.

**Figure 4:**
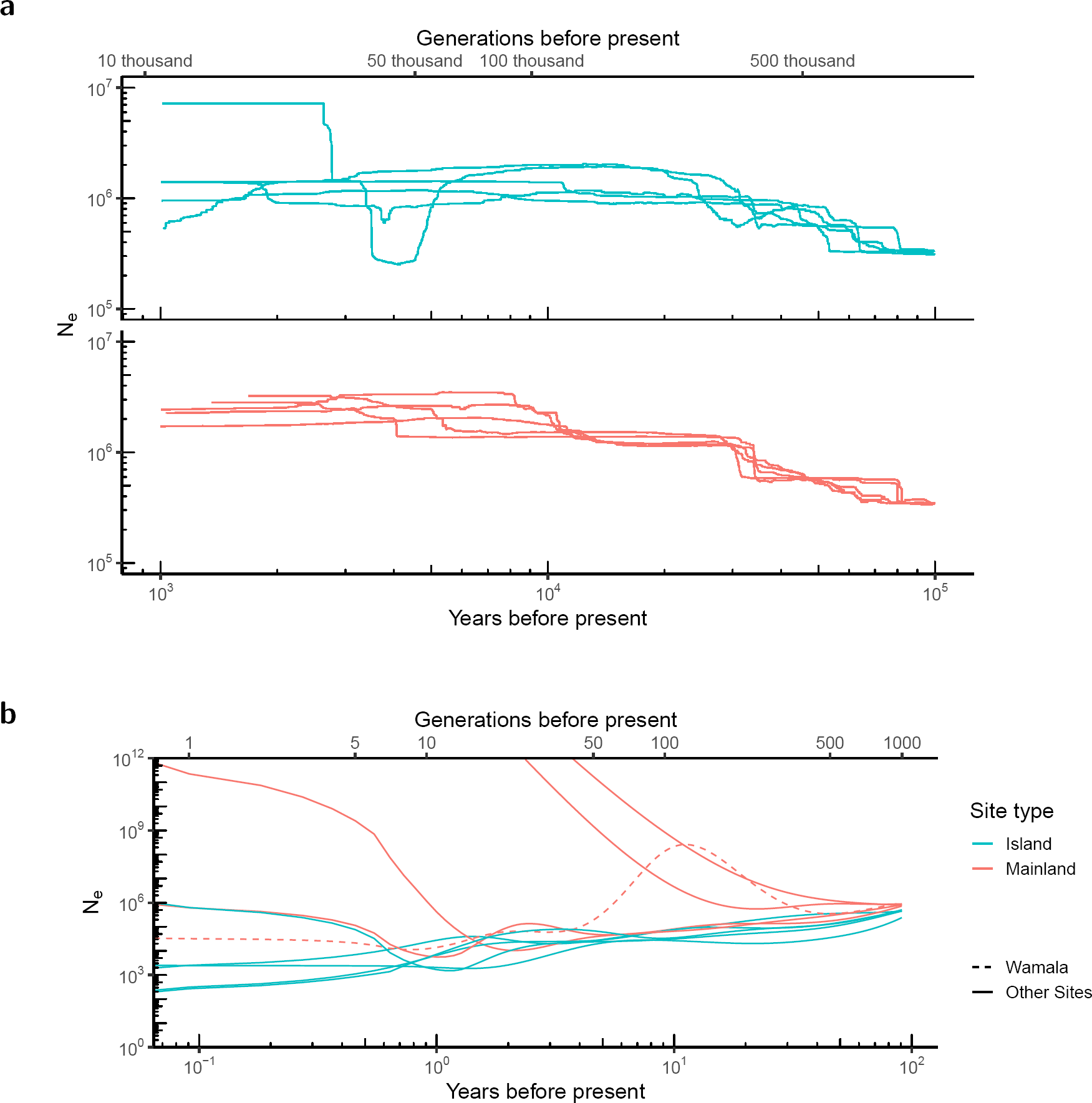
Population history of the Lake Victoria Basin samples. (A) Long-term evolutionary population histories inferred via stairway plots for island and mainland samples. (B) Contemporary or short-term effective population size (*N*_*e*_) history inferred using sharing of regions that are identical by descent (IBD). Wamala, a mainland locality showing island-like fluctuations in population size, is indicated with a dashed line. Plot truncated to exclude implausibly high estimates that are likely an artifact of sample size.

To all pairs of LVB localities we fit an isolation-with-migration (IM) demographic model using *δ*a*δ*i, in which an ancestral population splits into two populations, allowing exponential growth and continuous asymmetrical migration between the daughter populations (Supplementary Figs. S6, S7). In all comparisons involving islands and some between mainland sites, the best fitting model as chosen via AIC had zero migration (Supplementary Tables S3, S4, and S5). Time since population split was much more recent for mainland-mainland comparisons (excluding Bugala, median: 361 years) than those involving islands (island-island median: 7,128 years; island-mainland median: 4,043 years). Island-island split time confidence intervals typically did not overlap those involving mainland sites.

## Selection

As beneficial variants would be the most likely signatures of past gene flow to persist, we next examined signatures of selective sweeps for insight into migration. Identifying signatures of selection in the same genomic region in populations with independent lineages would be consistent with several scenarios [23]: (*i*) independent parallel selective sweeps on *de novo* mutations, (*ii*) independent parallel selective sweeps on shared ancestral variation, or (*iii*) selective sweeps on variants transferred via gene flow. As we were most interested in the transfer of adaptive variants for its insight into migration (*iii*), we distinguished between the alternative scenarios as follows.

We would expect independent sweeps on novel mutations (*i*) to exhibit differences in genetic background between the two populations, evidenced by distinct haplotype clusters, each comprising near-identical haplotypes separated by individual haplotypes lacking signatures of a selective sweep. In both other scenarios (*ii-iii*), we would instead expect haplotypes with the sweep to group together when clustered by genetic distance. By itself, haplotype information does not differentiate sweeps targeting standing ancestral variation (*ii*) from those targeting adaptive variants spread through gene flow (*iii*), but additional information such as geographic distance between the populations, estimates of gene flow inferred from other regions of the genome, and assessment of gene flow between other nearby populations, may suggest that one of these scenarios is the more likely.

While the sharing of a sweep may indicate migration between populations, the inverse would be suggestive—though not conclusive—of barriers to gene flow. A lack of sharing of a selective sweep signal between two populations may indicate no migration is occurring. However, it would also be consistent with the occurrence of migration that is subsequently countered by the local effects of selection or lost to genetic drift.

We first compared mainland Uganda and the Ssese Islands, reasoning that shared signatures of sweeps at a genomic location may indicate migration is occurring with the islands, while the absence was suggestive of isolation. We identified sweeps in the LVB using genome scans of between- and within-locality statistics, including *F*_*ST*_ ([24], Supplementary Fig. S8), Extended Haplotype Homozygosity (XP-EHH, [25], Supplementary Fig. S8), and haplotype homozygosity (H12, [26], Supplementary Fig. S9). To test for sweeps that were variable within the LVB, we identified locality-specific sweeps (found at only one sampling site in the LVB), sweeps that were found in our island localities but not mainland LVB localities, and sweeps that were found only in our mainland LVB localities (all defined as H12 *>* 99th percentile). To add additional country-level context, we then intersected these regions with those under putative selection in a mainland Ugandan reference population (H12 *>* 95th percentile; [4]).

Some genomic locations had heterogeneous selection signals within the LVB and within Uganda, indicative of potential geographic barriers to gene flow or local variation in selective regimes. Locality-specific putative sweeps were more prevalent on island than LVB mainland localities (mean per locality: island = 52.4; mainland = 26.8), concordant with increased isolation of the islands (Supplementary Table S6). Sweeps detected only or primarily in mainland LVB localities were shared with the Ag1000G Ugandan reference population more often (8 of 37; 22%) than those found only or primarily on islands (1 of 21, 5%; Supplementary Tables S7 and S8), again indicative of some barriers to gene flow with the islands.

We next reasoned that continent-wide selective sweeps, with broadly distributed selective advantage, would be the most likely to be shared via gene flow. Widespread sweeps that were absent or at extremely low frequency on the islands would be a strong suggestion against contemporary gene flow, and those that conversely were present on the islands would be indicative that gene flow had occurred, if the alternative scenarios could be excluded as outlined above. To identify these regions, we intersected our set of sweeps with those under putative selection in populations across the continent (H12 *>* 95th percentile in Ag1000G; [4]).

As expected, outlier regions included known selective sweep targets from elsewhere in Africa ([4], Supplementary Table S9). All sweeps found in the reference Uganda population [4] were detected in at least some sampling localities in our LVB dataset, except the sweep targeting *Vgsc*, which was likely excluded during filtration of the region adjacent to the centromere. For instance, the large genomic region spanning the cluster of insecticide resistance-associated cytochrome P450s (*Cyp6p*) on chromosome arm 2R, including *Cyp6p3* which is upregulated in mosquitoes with permethrin and bendiocarb resistance [27], exhibited low diversity, an excess of low frequency polymorphisms (Tajima’s *D*), and elevated haplotype homozygosity (H12) within the LVB populations (Supplementary Figs. S9, S10). Pairwise statistics (*F*_*ST*_ and XP-EHH) indicated low differentiation between LVB localities, as expected for a continent-wide sweep (Supplementary Figs. S8). The signal was found in every LVB site, including all islands. Hierarchical clustering of LVB and Ag1000G haplotypes revealed clades with low inter-haplotype diversity, expected after selection rapidly increases the frequency of a haplotype containing adaptive variation (Supplementary Fig. S11). Consistent with previous results [4], these clusters of closely related haplotypes on independent lineages indicate that multiple parallel sweeps targeting the *Cyp6p* region have occurred in several genetic backgrounds at numerous localities across Africa. Within Uganda, since almost all mainland and island individuals carry haplotypes from a single cluster, the selected haplotype of this cluster likely spread to near-fixation via gene flow.

In contrast, some sweeps with continent-wide prevalence including the reference Ugandan population [4] were found at all mainland LVB sites but had colonized the islands incompletely. For example, a region on chromosome arm 2L (2L:2,900,000-3,000,000) was found in all assayed Ag1000G populations and LVB mainland sites, but found on no island but Sserinya (Supplementary Table S8). As in previous studies [4], independent clusters of low-diversity haplotypes in varied genetic backgrounds suggest multiple sweeps targeting the cluster of genes encoding glutathione S-transferases (*Gste1* -*Gste7*), including one sweep specific to Uganda. This Ugandan sweep was similarly confined largely to the mainland in the LVB. These sweeps at targets of selection throughout the continent that are largely restricted to the mainland are suggestive of strong barriers to gene flow to the islands, either due to lack of connectivity or the countering effects of selection or drift. Other sweeps had colonized the islands incompletely. The sweep targeting cytochrome P450 gene *Cyp9k1* likely arose multiple times independently, since Ugandan haplotypes do not cluster with low diversity clusters from elsewhere in Africa. Within the LVB, the sweep signature is found on some, but not all islands, suggesting some barrier to gene flow or local selection limiting the spread of the sweep.

Two regions exhibited selection signals similar in amplitude to known insecticide-related loci, with elevated between-locality differentiation, low diversity, and extended homozygosity (Supplementary Figs. S8, S9, S12, and S13). The first, at 2L:34.1 Mb, contains many genes, including a cluster involved in chorion formation [28] near the signal peak. Haplotype clustering revealed a group of closely-related Ugandan individuals, consistent with a geographically bounded selective sweep (Supplementary Fig. S14). The selected variation had not fully colonized the islands or the LVB mainland sites, however, suggesting some barriers to gene flow, loss due to drift at some localities, or local differences in selective pressure within the LVB. Elsewhere in Africa, clustering analysis revealed other low-variation clades in distinct genetic backgrounds in, *e.g.*, Cameroon and Angola, suggesting parallel selection on independent mutations at this locus.

The second putative sweep, at X:9.2 Mb, coincided precisely with eye-specific diacylglycerol kinase (AGAP000519, chrX:9,215,505-9,266,532). Low diversity haplotypes formed a single cluster including LVB haplotypes overwhelmingly from the islands (Supplementary Fig. S15). Transfer via gene flow between islands but not to the mainland is reasonable, given the connectivity patterns we have inferred from neutral variation. Additionally, local selection may be countering the spread of the sweep to the mainland. However, more surprisingly, these island haplotypes with evidence of a selective sweep were most closely related to haplotypes from distant locations, primarily Gabon and Burkina Faso rather than Uganda. This sharing of extended haplotypes between islands and distant localities is consistent with either gene flow or independent sweeps targeting ancestral standing variation. Of these alternatives, extremely long distance gene flow that persists only on islands seems less likely.

## Discussion

Understanding the population genetics of island *Anopheles gambiae* has both evolutionary and practical importance. A limited number of genetic investigations have been conducted on oceanic [29–32] and lacustrine islands [33–36], though the latter have been limited in the type or count of molecular markers used. In contrast to shallow population structure across Africa [4, 5], partitioning of genetic variation on islands suggests varying isolation. Using a genome-wide dataset, we found differentiation between the Ssese Islands to be relatively high in the context of continent-wide structure, with the differentiation between Banda Island (only 30 km offshore) and mainland localities on par with or higher than for populations on opposite sides of the continent (*e.g.*, Banda vs. Wamala, *F*_*ST*_ = 0.034; mainland Uganda vs. Burkina Faso, *F*_*ST*_ = 0.007 [4]). The Ssese Islands are approximately as differentiated as all but the most outlying oceanic islands tested (*e.g.* mainland Tanzania vs. Comoros, 690-830 km apart, *F*_*ST*_ = 0.199-0.250 [31]). Patterns of haplotype sharing did include direct evidence for the recent exchange of migrants between nearby islands, but analyses based on haplotype sharing, Bayesian clustering, and demographic reconstruction included no evidence of direct sharing between Banda and the mainland. Banda is nonetheless connected to other islands and thereby indirectly connected to the mainland, and additional sampling may reveal signs of admixture. Additional sampling on Banda and other islands that are disjunct from the rest of the archipelago would be prudent when assessing potential field testing locations.

The name “Ssese” derives from another arthropod vector, the tsetse fly (*Glossina* spp.) The tsetse-mediated arrival of sleeping sickness in 1902 brought “enormous mortality” [37, pp. 332] to the 20 thousand residents, who were evacuated in 1909 [37, 38]. Though encouraged to return by 1920, the human population numbered only 4 thousand in 1941 [37] and took until 1980 to double [39], but has since rapidly risen to over 62 thousand (2015, projected; [18, 40]). The impacts on mosquito populations of this prolonged depression in human population size, coupled with water barriers to mosquito migration, are reflected in the distinctive demographic histories of island *An. gambiae* populations, which were smaller and fluctuated more than mainland localities, echoing previous results [34, 36]. Two mainland sites had island-like recent population histories, with Wamala abruptly switching from a mainland-like to island-like growth pattern around 2005. This coincides precisely with a *≥* 20% reduction from 2000-2010 in the *Plasmodium falciparum* parasite rate (P*f* PR_2−10_; a measure of malaria transmission intensity) in Mityana, the district containing Wamala [41].

Though previous *Anopheles* population genetic studies have inferred gene flow even among species [4, 42], we inferred that no genetic exchange had occurred since the split between island sites and between islands and the mainland. Island pairs were inferred to have split far deeper in the past (5,000-14,000 years ago) than mainland sites (typically *<* 500 years ago), on par with the inferred split time between Uganda and Kenya (approximately 4,000 years ago; [4]). Although bootstrapping-derived confidence intervals permit some certainty, our model fit is not optimal likely due to low sample sizes and high levels of shared ancestral variation, and additional sampling is necessary to clarify population history. Our inferred lack of gene flow to the islands appears contradictory to the presence of individuals who share ancestry with the mainland on all islands but Banda. We cannot dismiss the possibility that this indicates actual migration occurs. If so, effects of migration would have to be sufficiently countered by local selection to limit its effect on allele frequency spectra, rendering effective migration (as estimated in population history inference) zero. The apparent contradiction can also be resolved if shared ancestry between islands and mainland suggested by the clustering result is interpreted as retention of shared ancestral polymorphism or the existence of inadequately sampled ancestral variation [43], rather than recent admixture. This interpretation is consistent with the affinity we observed between the Ssese Islands and West Africa in the structure of adaptive variation.

Discerning whether the absence of observed gene flow is due to lack of connectivity, the opposition of selection, or the stochasticity of genetic drift is difficult. Instead we must rely on estimates of the strength of selection in the two locales to inform our conclusions. For example, we would expect that an insecticide sweep found all over Africa would spread in island mosquito populations with insecticide treated bed nets, despite the considerable effect of genetic drift in small populations. As insecticide treated bed net usage is present on the islands [18], variation conferring a major selective advantage related to insecticides would be expected to spread to and persist on the islands if migration allows the transfer, and the strongest evidence of a lack of contemporary connectivity is therefore the absence of a sweep on the islands that is widespread on the continent.

We found two sweeps on insecticide-related genes that are common targets of selection elsewhere but which have incompletely colonized the Ssese Islands: one on cytochrome P450 monooxygenase *Cyp9K1* [44, 45] present on some islands, and another on glutathione Stransferase genes (*Gste1-Gste7*; [46–49]) at extremely low frequency on the islands. That the selective sweeps targeting these loci [4] have not fully colonized the islands despite the advantage in detoxifying pyrethroids and DDT suggests a lack of contemporary exchange. However, the sweep targeting the *Cyp6p* cluster was found on all islands, confirming past gene flow had occurred at some point. Although these distributions confirm that past migration from the mainland to islands has occurred and we are unable to exclude low levels of contemporary gene flow, taken together our data are consistent with potentially high degrees of isolation on contemporary timescales for some islands of the Ssese archipelago.

Our investigation also identified two previously unknown signatures of selection. For the first, on chromosome arm 2L and encompassing many genes, haplotypes with sweeps in distinct genetic backgrounds across Africa suggest the region has been affected by multiple independent convergent sweeps. In Uganda, most individuals with the sweep are from the mainland, suggesting a local origin and spread via short distance migration. The putative target of the second sweep is diacylglycerol kinase on the X-chromosome, a homolog of retinal degeneration A (*rdgA*) in *Drosophila*. The gene is highly pleiotropic, contributing to signal transduction in the fly visual system [50, 51], but also olfactory [52] and auditory [53] sensory processing. It has been recently implicated in nutritional homeostasis in *Drosophila* [54] and is known to interact with the TOR pathway [55], which has been identified as a target of ecological adaptation in *Drosophila* [56, 57] and *An. gambiae* [58]. The sweep appears largely confined to island individuals in the LVB, but the cluster of haplotypes also includes those from Gabon, Burkina Faso, and Kenya. Shared extended haplotypes suggest a single sweep event spread by gene flow or selection on standing ancestral variation, not independent selection on *de novo* mutations. Possible explanations include long distance migration of an adaptive variant persisting on only the islands or, more reasonably, selection on standing ancestral variation. We have not found obvious candidate targets of selection, *e.g.* coding changes, which may be due to imperfect annotation of the genome or the likely possibility that the target is a non-coding regulator of transcription or was filtered from our dataset. Further functional studies would be needed to clarify the selective advantage that these haplotypes confer. Interestingly, the putative sweep coincides with a similar region of low diversity in a cryptic subgroup of *Anopheles gambiae sensu lato* (GOUNDRY; [42]), suggesting possible parallel selective events on independent mutations or adaptive introgression.

Population structure investigations are paramount for informing the design and deployment of control strategies, including field trials of transgenic mosquitoes. We demonstrate alternatives to simple extrapolation of migration rates from differentiation, which is fraught [59] particularly given the assumption of equilibrium between the evolutionary forces of migration and drift [59–61], an unlikely state for huge *An. gambiae* populations [3]. We suggest that future assessments of connectivity include, as we have, the spatial distribution of adaptive variation, identification of recent migrants via haplotype sharing, and demographic history modeling, from which we have inferred the Ssese Islands to be relatively isolated on contemporary time scales. Though we cannot exclude the possibility of a small amount of gene flow over evolutionary time between our most isolated islands and the mainland, the data are consistent with a sufficiently low amount of gene flow that it becomes reasonable to consider these islands as isolated on short time frames.

A completely isolated population of mosquitoes is not a reasonable expectation given mosquitoes’ propensity for active and even passive (human-aided or windborne) dispersal [16], potentially up to hundreds of kilometers [11]. Although no island, lacustrine or oceanic, is completely isolated, such localities may still be ideal for initial gene drive field testing, as the geographical barriers maximize isolation to the extent possible [16], and absolute isolation on evolutionary timescales is unnecessary given the relatively short timeframe of small-scale field tests. Thus, the probability of contemporary migration may be sufficiently low to qualify some Ssese Islands as candidate field sites. Additionally, the assessment of the islands’ suitability as potential sites for field trials of genetically modified mosquitoes must also consider the logistical ease of access and monitoring that the bounded geography of a small lacustrine island with low human population density affords initial field tests. Due consideration should be provided to these characteristics of small lake islands that may be appealing to regulators, field scientists, local communities, and other stakeholders. Given such features and the probable rarity of migration, the Ssese Islands may be logical and tractable candidates for initial field tests of genetically modified *An. gambiae* mosquitoes, warranting further entomological study.

## Materials and Methods

### Experimental design

Mosquitoes were sampled from 5 of the Ssese Islands in Lake Victoria, Uganda (Banda, Bukasa, Bugala, Nsadzi, and Sserinya) and 4 mainland sampling localities (Buwama, Kaazi, Kiyindi, and Wamala) at varying distances from the lake in May and June, 2015. Sampling took place between 4:40 and 8:15 over a 30 day period as follows: Indoor resting mosquitoes were collected from residences via mouth or mechanical aspirators and subsequently identified morphologically to species group. Female mosquitoes assigned to the *An. gambiae sensu lato* complex based on morphology (*N* =575) were included in further analyses. All mosquitoes were preserved with silica desiccant and transported to the University of Notre Dame, Indiana, U.S.A. for analysis.

### DNA extraction, **Library preparation**, **and Whole Genome Sequencing**

Animals were assigned to species level via a PCR-based assay [62] using DNA present in a single leg or wing. DNA from individual *An. gambiae s. s.* N=116 mosquitoes was extracted from the whole body via phenol-chloroform extraction [63] and then quantified via fluorometry (PicoGreen). Automated library preparation took place at the NYU Langone Medical Center with the Biomek SPRIWorks HT system using KAPA Library Preparation Kits, and libraries were sequenced on the Illumina HiSeq 2500 with 100 paired end cycles.

### Mapping and SNP calling, **filtering**

Software version information is provided in Supplementary Table S10. After quality filtering and trimming using ea-utils’ fastq-mcf (-l 15 -q 15 -w 4; [64]), reads were mapped to the *An. gambiae* reference genome (AgamP4 PEST; [65, 66]) using BWA aln and sampe with default parameters [67].

After realignment around indels with GATK’s IndelRealigner, variants were called using GATK’s UnifiedGenotyper (with -stand call conf 50.0 and -stand emit conf 10.0; selected to be consistent with methods of recent comparison SNP dataset [4]) and filtered for quality [68], excluding SNPs with QualByDepth (QD) *<* 2.0, RMSMappingQuality (MQ) *<* 40.0, FisherStrand (FS) *>* 60.0, HaplotypeScore *>* 13.0, or ReadPosRankSum *<* −8.0. All bioinformatic steps for read mapping and variant identification are encapsulated in the NGS-map pipeline (https://github.com/bergeycm/NGS-map). This yielded 33.1 million SNPs. Individuals and variants with high levels of missingness (*>* 10%) and variants that were not biallelic or exhibited values of HWE that were likely due to sequencing error (p *<* 0.00001) were excluded from further analysis. For use in population structure inference, the SNP dataset was further pruned for linkage disequilibrium by sliding a window of 50 SNPs across the genome in 5 SNP increments and recursively removing random SNPs in pairs with *r*^2^ *>* 0.5 using PLINK [69, 70]. After filtration, the dataset contained 28,569,621 SNPs before LD pruning and 115 individuals. SNPs unpruned for linkage disequilibrium were phased with SHAPEIT2 [71] using an effective population size (*N*_*e*_) of 1,000,000 (consistent with previous demographic modeling [4]), default MCMC parameters (7 burn-in MCMC iterations, 8 pruning iterations, and 20 main iterations), conditioning states for haplotype estimation (*K* = 100), and window size of 2 Mb.

### Population structure inference

To explore population structure in a larger, continentwide context, we merged our LVB SNP set with a recently published dataset of *Anopheles gambiae* individuals (from the Ag1000G project) [4]. Prior to filtering, biallelic SNPs from the LVB and Ag1000G datasets were merged using bcftools [72]. We excluded any SNP with greater than 10% missingness in either dataset, any SNPs that did not pass the accessibility filter of the Ag1000G dataset, and SNPs with MAF *<* 1%. After this filtration, our merged SNP dataset contained 12,537,007 SNPs.

After pruning the merged dataset for LD (leaving 9,861,756 SNPs) and excluding laboratory crosses (leaving 881 individuals), we assigned individuals’ genomes to ancestry components using ADMIXTURE [20]. We created 10 replicate samples of 100,000 SNPs from chromosome 3 (prior to LD-pruning), including only biallelic SNPs in euchromatic regions with MAF *>* 1%. These replicate datasets were pruned for LD by randomly selecting from pairs of SNPs with *r*^2^ *>* 0.01 in sliding windows of size 500 SNPs and with a stepsize of 250 SNPs. For each replicate, we ran ADMIXTURE for 5 iterations in five-fold cross validation mode for values of *k* from 2 to 10. This resulted in 50 estimates for each value of *k*. We assessed these results using the online version of CLUMPAK with default settings to ensure the stability of the resulting clustering [73]. CLUMPAK clusters the replicate runs’ Q-matrices to produce a major cluster for each value of *k*, which we then visualized. The lowest cross-validation error was found for *k* = 6 clusters, but we also display ancestry estimates with *k* = 9 clusters to further explore patterns of structure with a level of subdivision at which the Ssese Island individuals are assigned a unique ancestry component.

We visualized population structure via principal components analysis (PCA) with PLINK [69, 70], using the LVB-Ag1000G merged dataset (excluding the outlier Kenyan population; [4]) and 3,212,485 chromosome 3 SNPs (to avoid the well-known inversions on chromosome 2 and the X-chromosome) outside of heterochromatic regions (such as centromeric regions; [66]; Supplementary Table S11). We next performed a PCA on the LVB dataset alone, pruning for LD and low-MAF (*<* 1%) SNPs on chromosome 3. Based on the results of this analyses, we split individuals from the large island of Bugala into two clusters for subsequent analyses: those that cluster with mainland individuals and those that cluster with individuals from the smaller islands.

We computed the pairwise fixation index (*F*_*ST*_) between locality samples for *An. gambiae* using the unbiased estimator of Hudson [74] as implemented in smartpca [75, 76]. To obtain overall values between sampling sites, per-SNP values were averaged across the genome excluding known inversions (2*La*, 2*Rb*, and 2*Rc*) and heterochromatic regions. We also computed *z*-scores via block jackknife, using 42 blocks of size 5 Mb. We tested for isolation by distance, or a correlation between genetic and geographic distances, with a Mantel test [77] as implemented in the R package ade4 [78], using these *F*_*ST*_ estimates and Euclidean geographic distances between localities.

To estimate fine-scale structure and relatedness between individuals, we estimated the proportion of pairs of individuals genomes that are identical by descent (IBD) using PLINK [69, 70]. We excluded heterochromatic and inversion regions, and retained informative pairs of SNPs within 500 kb in the pairwise population concordance test.

### Diversity estimation

Grouping individuals by site (except for Bugala, which was split based on the results of the PCA), we calculated nucleotide diversity (*π*) and Tajima’s *D* in nonoverlapping windows of size 10 kb, the inbreeding coefficient (*F*) estimated with the method of moments, minor allele frequencies (the site frequency spectrum, SFS), and a measure of linkage disequilibrium (*r*^2^) using VCFtools (Danecek2011). For *r*^2^, we computed the measure for all SNPs (unpruned for linkage) within 50 kb of a random set of 100 SNPs with MAF *>* 10% and corrected for differences in sample size by subtracting 1*/n*, where *n* equaled the number of sampled chromosomes per site. To visualize decay in LD, we plotted *r*^2^ between SNPs against their physical distance in base pairs, first smoothing the data by fitting a generalized additive model (GAM) to them. We also inferred runs of homozygosity using PLINK [69, 70] to compare their length (*F*_*ROH*_), requiring 10 homozygous SNPs spanning a distance of 100 kb and allowing for 3 heterozygous and 5 missing SNPs in the window. Runs of homozygosity were inferred using LD-pruned SNPs outside of inversions or heterochromatic regions. We tested the significance of differences in these statistics between island and mainland categories using a two-sided Wilcoxon rank sum test.

### Demographic history inference

To estimate the contemporary or short-term *N*_*e*_ for each site, we inferred regions of IBD from unphased data with IBDseq [79] and analyzed them with IBDNe [22]. We restricted our analysis to SNPs from chromosome 3 to avoid inverted regions. We allowed a minimum IBD tract length of 0.005 cM (or 5 kb), scaling it down from the recommended length for human genomes due to mosquitoes’ high level of heterozygosity [4] and assumed a constant recombination rate of 2.0 cM/Mb (after [80]).

To estimate the long-term evolutionary demographic history of mosquitoes on and near the Ssese Islands, including a long-term estimate of *N*_*e*_ [81], we inferred population demographic history for each site via stairway plots using the full site frequency spectra based on SNPs on chromosome 3 with heterochromatic regions and regions within 5 kb of a gene excluded [21].

We also inferred a “two-population” isolation-with-migration (IM) demographic model with *δ*a*δ*i [82, 83] in which the ancestral population splits to form two daughter populations that are allowed to grow exponentially and exchange migrants asymmetrically, as described in the main text. For *δ*a*δ*i-based analyses, we used the full dataset of SNPs on chromosome 3, not pruned for LD but with heterochromatic regions and regions within 5 kb of a gene masked. We polarized the SNPs using outgroup information from *Anopheles merus* and *An. merus* [84]. We fit this two-population model and the same model without migration to all pairs of locality samples, choosing the optimal model using the Godambe Information Matrix and an adjusted likelihood ratio test to compare the two nested models. We compared the test statistic to a *χ*^2^ distribution and rejected the null model if the p-value for the test statistic was *<* 0.05. For both, singletons and doubletons private to one population were masked from the analysis and a parameter encompassing genotype uncertainty was included in the models and found to be low (mean = 0.0.70%). We assessed the goodness-of-fit visually using the residuals of the comparison between model and data frequency spectra (Supplementary Fig. S7). Using the site frequency spectrum, we projected down to 2-6 fewer chromosomes than the total for the smaller population to maximize information given missing data. We set the grid points to {*n, n* + 10, *n* + 20}, where *n* = the number of chromosomes. Bounds for *N*_*e*_ scalars were *ν ∈* (0.01, 10, 000), for time were *T ∈* (1*e*-8, 0.1), for migration were *m ∈* (1*e*8, 10), and for genotyping uncertainty were *p*_*misid*_ *∈* (1*e*-8, 1). Parameters were perturbed before allowing up to 1000 iterations for optimization. We estimated parameter uncertainty using the Fisher information matrix and 100 bootstrap replicates of 1 Mb from the dataset. If the Hessian was found to be not invertible when computing the Fisher information matrix, the results of that iteration were excluded from the analysis. For population size change parameters, *ν*, optimized values for one or both populations were often close to the upper limit. Due to this runaway behavior, common in analyses of the SFS [85], we excluded the population size change from our interpretation.

To translate *δ*a*δ*i- and stairway plot-based estimates of *N*_*e*_ and time to individuals and years respectively, we assumed a generation time of 11 per year and a mutation rate of 3.5*e*-9 per generation [4].

### Selection inference

To infer candidate genes and regions with selection histories that varied geographically, we compared allele frequencies and haplotype diversity between the sampling sites. To infer differing selection between sampling sites, we computed *F*_*ST*_ between all populations in windows of size 10 kb using the estimator of Weir and Cockerham [24] (as implemented in VCFtools [86]), and H12 (as implemented in SelectionHapStats [26]) and XP-EHH on a per-site basis (as implemented in selscan [87]) to detect long stretches of homozygosity in a given population considered alone or relative to another population [25]. For XP-EHH, EHH was calculated in windows of size 100 kb in each direction from core SNPs, allowing EHH decay curves to extend up to 1 Mb from the core, and SNPs with MAF *<* 0.05 were excluded from consideration as a core SNP. As we lacked a fine-scale genetic map for *Anopheles*, we assumed a constant recombination rate of 2.0 cM/Mb (after [80]). Scores were normalized within chromosomal arms and the X-chromosome. The between-locality statistics, *F*_*ST*_ and XP-EHH, were summarized using the composite selection score [CSS; [88, 89]].

We plotted these statistics across the genome to identify candidate regions with signatures of selection, including high differentiation between samples from different localities, reduced variability within a sample, and extended haplotype homozygosity. To identify regions of the genome showing signatures of selection specific to certain geographic areas, we identified genomic regions with elevated H12 in a subset of localities, and confirmed both elevated differentiation (as inferred from *F*_*ST*_) and evidence of differing selective sweep histories (as inferred from XP-EHH). Excluding the mainland-like portion of Bugala, we identified putative locality-specific sweeps (H12 over 99^th^ percentile in one population), island-specific sweeps (H12 over 99^th^ percentile in 4 or more of the 5 island localities but 0 or 1 mainland localities), or LVB mainland-specific sweeps (H12 over 99^th^ percentile in 3 or more of the 4 island localities but 0 or 1 island localities). To place these putative sweeps in their continental context, for the region of each putative locality-, island-, or LVB mainland-specific sweep, we determined if the H12 values of each of the Ag1000G populations (excluding Kenya due to its signatures of admixture and recent population decline; [4]) were in the top 5% for that population, indicating a possible selective sweep at the same location.

We further explored the haplotype structure and putative functional impact of loci for which we detected signatures of potential selection to determine the count and geographic distribution of independent selective sweeps. To provide necessary context for the reconstruction of sweeps and quantify long distance haplotype sharing between populations, we included data from several other *An. gambiae* populations across Africa (Burkina Faso, Cameroon, Gabon, Guinea, Guinea-Bissau, Kenya, and other Ugandan individuals; [4]). We computed the pairwise distance matrix as the raw number of base pairs that differed and grouped haplotypes via hierarchical clustering analysis (implemented in the hclust R function) in regions of size 100 kb centered on each peak in pairwise *F*_*ST*_ or XP-EHH, or the average of peaks, in the case for multiple nearby spikes. As short terminal branches can result from a beneficial allele and linked variants rising to fixation during a recent selective sweep, we identified such clusters by cutting the tree at a height of 0.4 SNP differences per kb.

## Acknowledgments

The authors would like to thank the UVRI field entomology team: Christine Babirye, Ronald Mayanja, Paul Mawejje, Kevin Nakato, and Fred Ssenfuka. We thank Nicholas Harding and Alistair Miles for helpful discussion. This study was supported by Target Malaria, which receives core funding from the Bill & Melinda Gates Foundation and from the Open Philanthropy Project Fund, an advised fund of Silicon Valley Community Foundation, through subcontracts to J.K.K. and N.J.B. N.J.B. also received support from NIH R01 AI125360 and R21 AI123491. The NYU School of Medicine’s Genome Technology Center is partially supported by the Cancer Center Support Grant P30CA016087 at the Laura and Isaac Perlmutter Cancer Center.

## Data and materials availability

All scripts used in the analysis are available at https://github.com/bergeycm/Anopheles gambiae structure LVB and released under the GNU General Public License v3. Sequencing read data for the LVB individuals are deposited in the NCBI Short Read Archive (SRA) under BioProject accession PRJNA493853.

## Author Contributions

C.M.B., J.K.K., and N.J.B. designed the study; C.M.B., M.L., R.M.W., and J.K.K. collected biological samples; C.M.B. analyzed the data; C.M.B., M.C.F., and N.J.B. wrote the manuscript; M.C.F., J.K.K., and N.J.B. supervised the research; C.M.B., M.L., R.M.W., M.C.F., J.K.K., and N.J.B. edited the manuscript.

## Supplemental Material for

**Figure S1:**
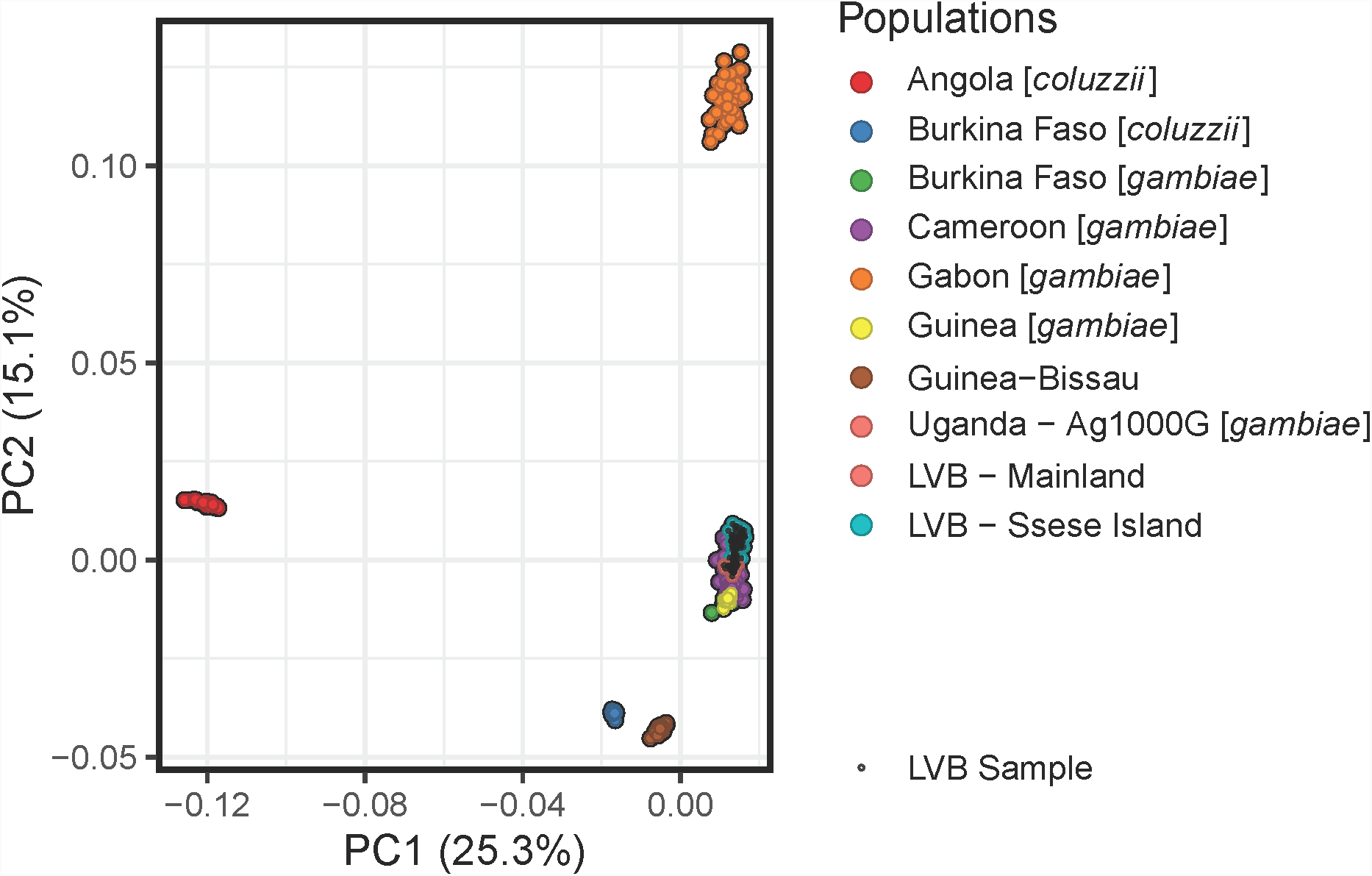
PCA plot of study individuals and *A. gambiae* and *A. coluzzii* individuals from reference Ag1000G populations, showing the first and second components. Kenyan population is not included, and analysis is based on chromosome 3.

**Figure S2:**
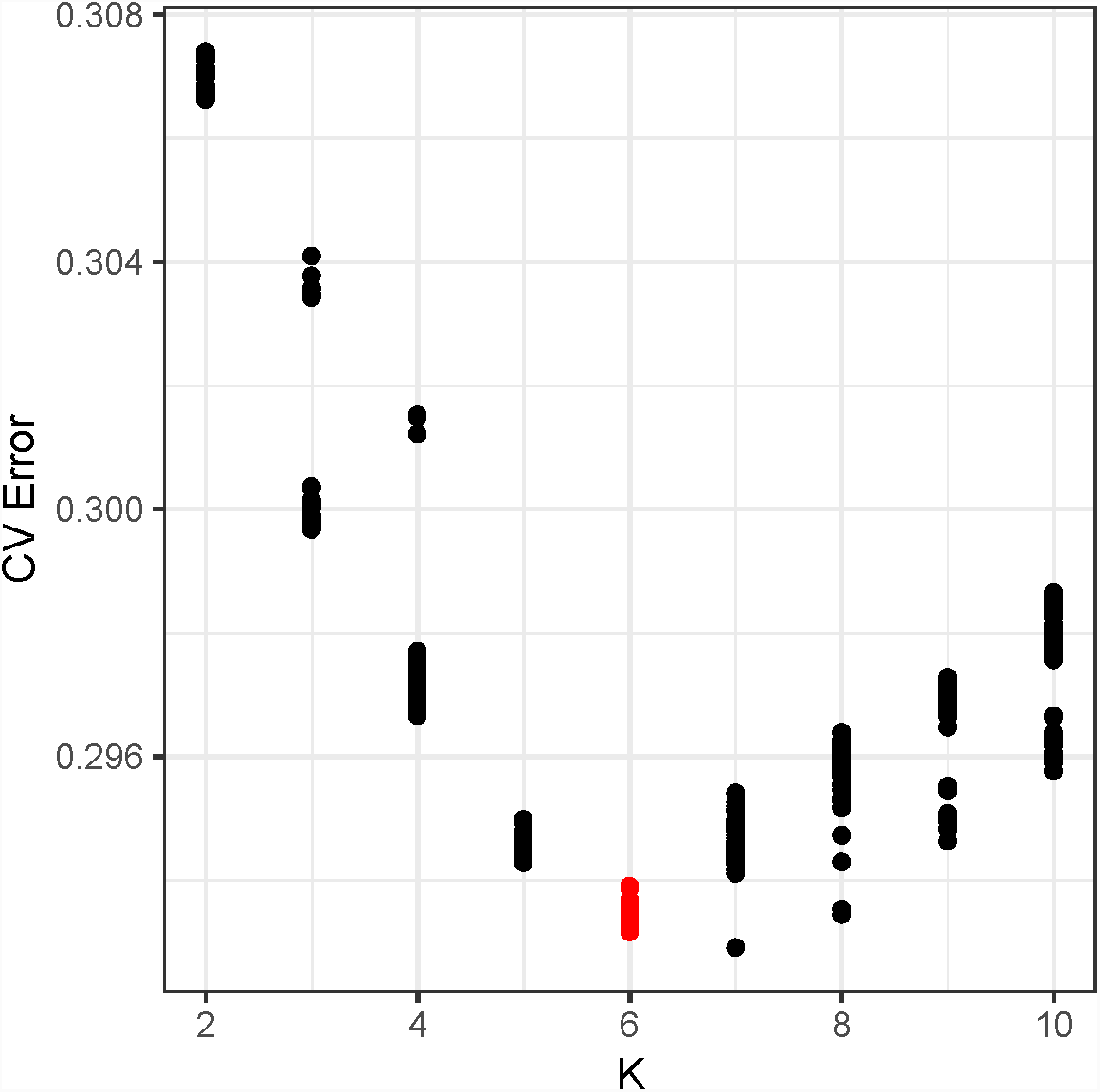
ADMIXTURE cross-validation error. Cross-validation error for range of *k* values for ADMIXTURE analysis of Lake Victoria Basin individuals and *A. gambiae* and *A. coluzzii* Ag1000G reference populations.

**Figure S3:**
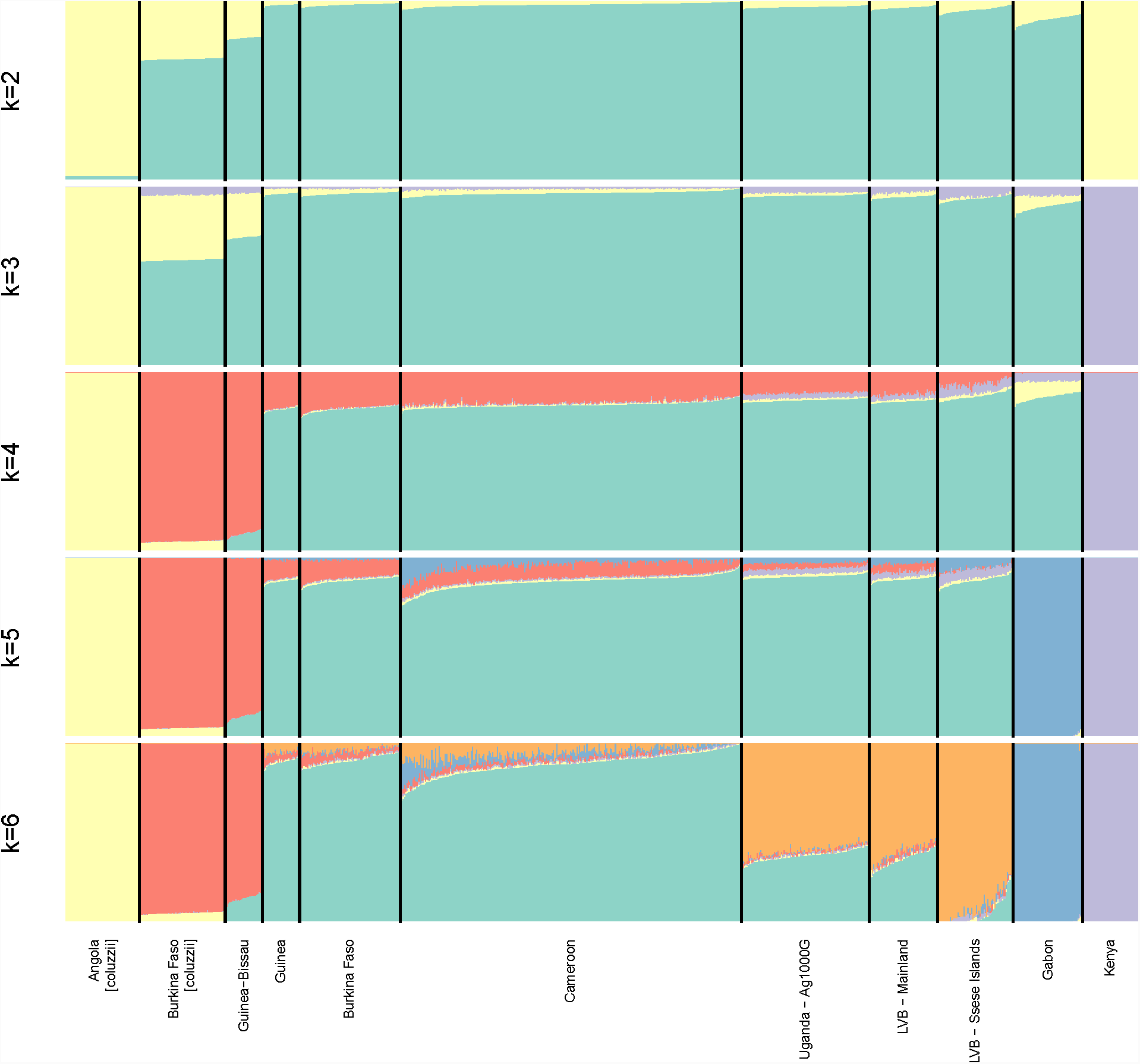
ADMIXTURE-inferred ancestry. Ancestry of individuals in Lake Victoria Basin and of Ag1000G reference populations as inferred by ADMIXTURE clustering method. Samples are *A. gambiae* unless noted, and analysis is based on chromosome 3. Using *k* = 6 clusters minimizes cross validation error (Fig. S2).

**Figure S4:**
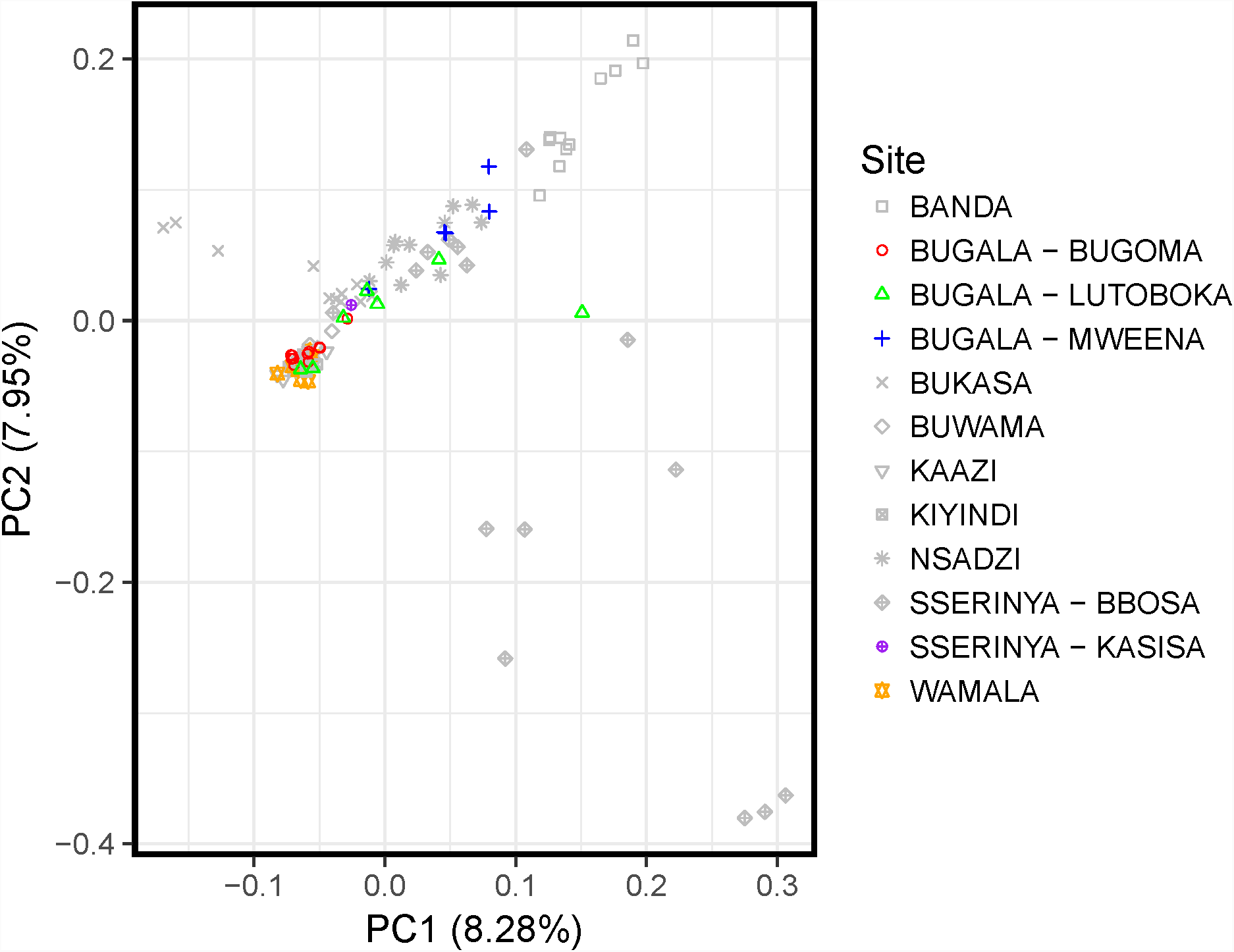
PCA showing Bugala subdivision. PCA colored by sampling locations. Based on this analysis, individuals from Bugala were split into mainland- and island-like subpopulations. Samples from Sserinya Island, though sampled from two localities, were not split.

**Figure S5:**
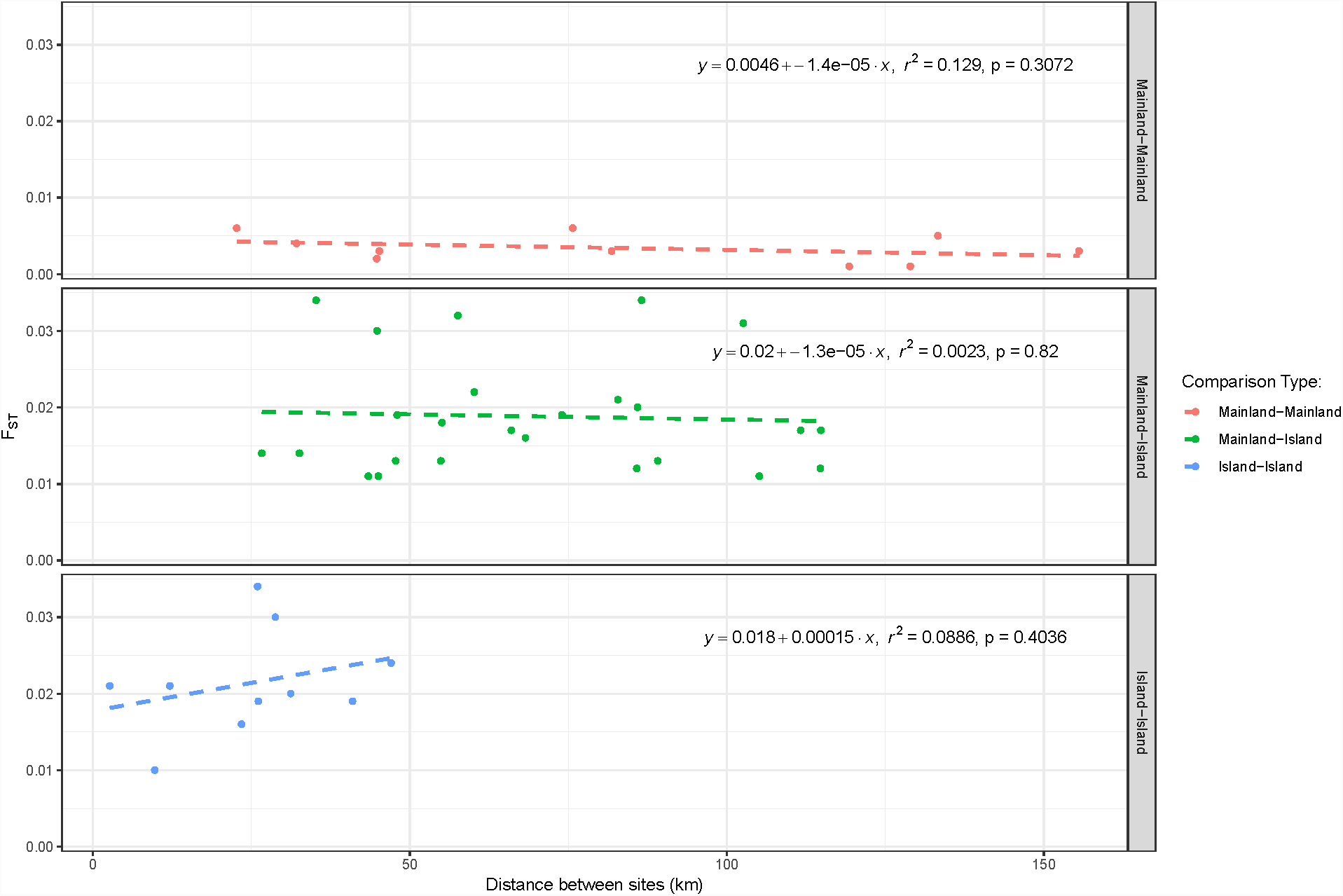
Correlations between genetic distance (*F*_*ST*_) and geographic distance between localities. The *p*-values are for the test that the slope is significantly different from zero.

**Figure S6:**
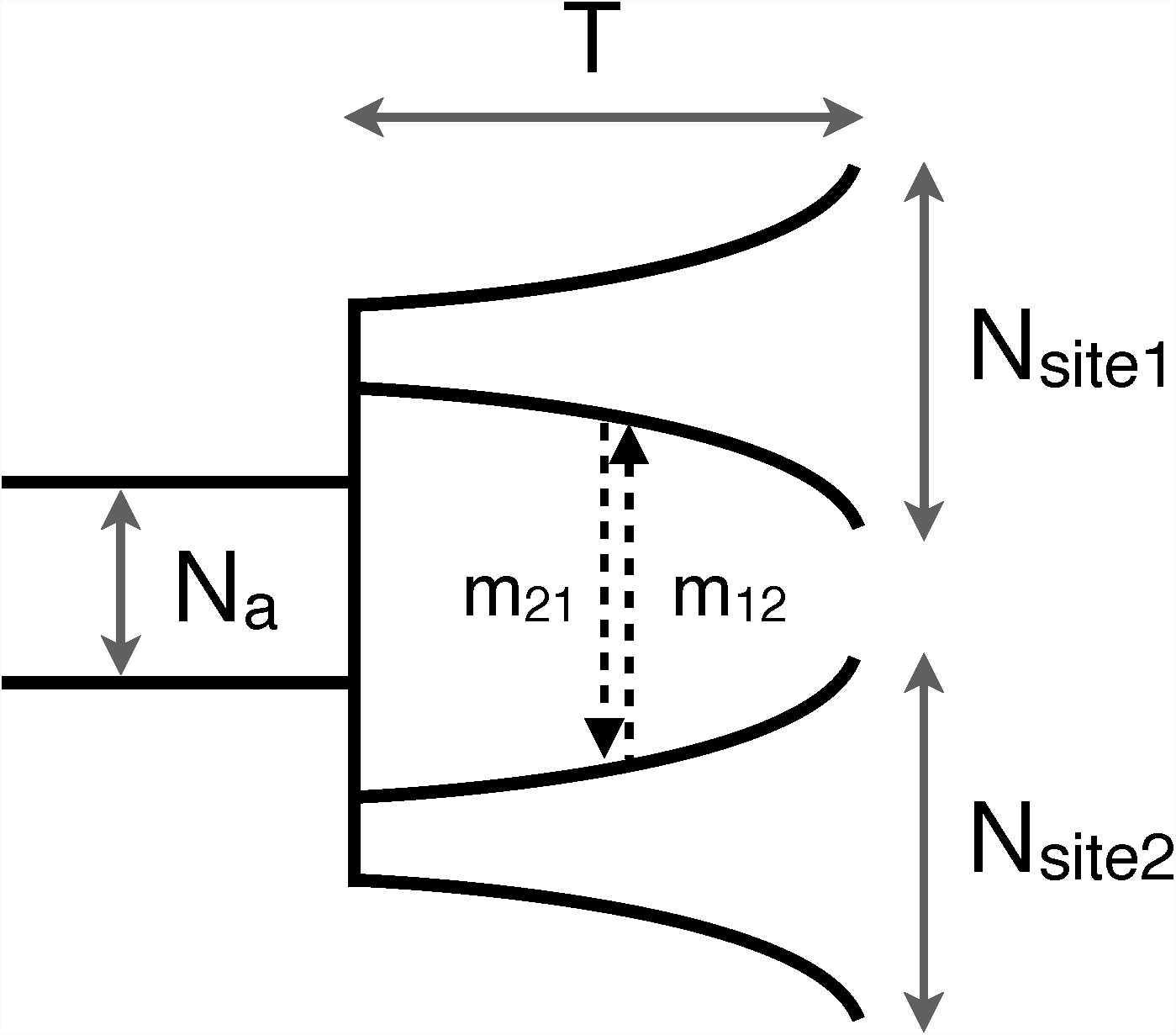
IM model schematics. Schematic of model fit to data with *δ*a*δ*i for population history inference between all pairs of sampled sites using IM model.

**Figure S7:**
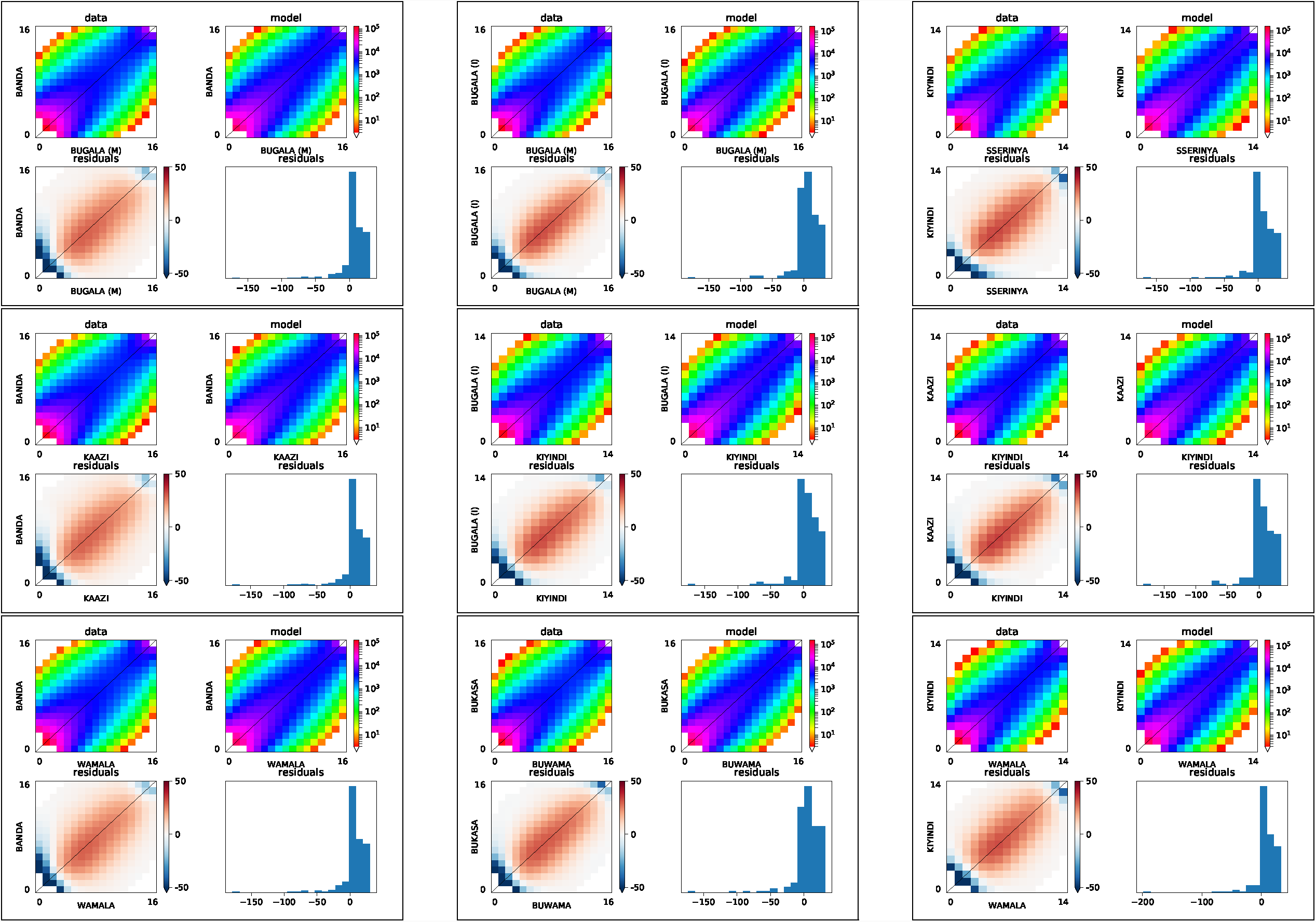
Two population *δ*a*δ*i optimization results. Comparison between best fitting model and data frequency spectra for two population *δ*a*δ*i inference. Of the pairwise comparisons for which the best model included migration, a randomly selected set of nine are shown here. Two-dimensional frequency spectra are plotted as logarithmic colormaps for the data (upper left) and model (upper right), and the bottom row plots show the residuals between model and data. Positive residuals in red indicate the model predicts too many SNPs in that entry while negative residuals in blue indicate the model predicts too few.

**Figure S8:**
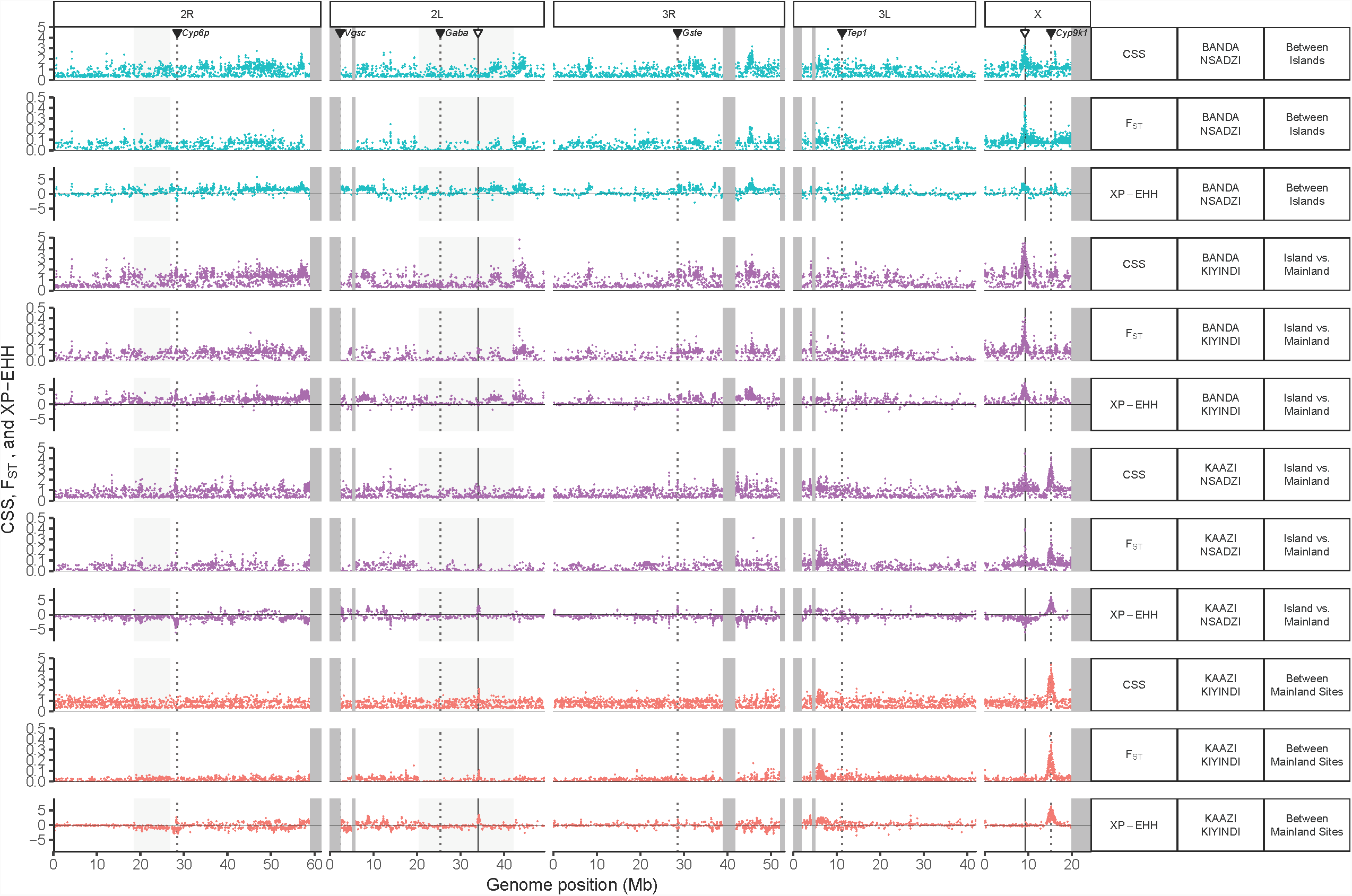
*F*_*ST*_, XP-EHH, and Composite Selection Score (CSS) across genome. *F*_*ST*_, XP-EHH, and CSS averaged in windows of size 10 kb plotted across genome for pairwise comparisons of island and mainland localities. Shaded regions indicate inversions or heterochromatic regions (excluded from analysis) and dotted lines indicate known insecticide genes while solid lines indicate the two putative sweeps identified in the present study. Only several exemplar pairs of populations shown.

**Figure S9:**
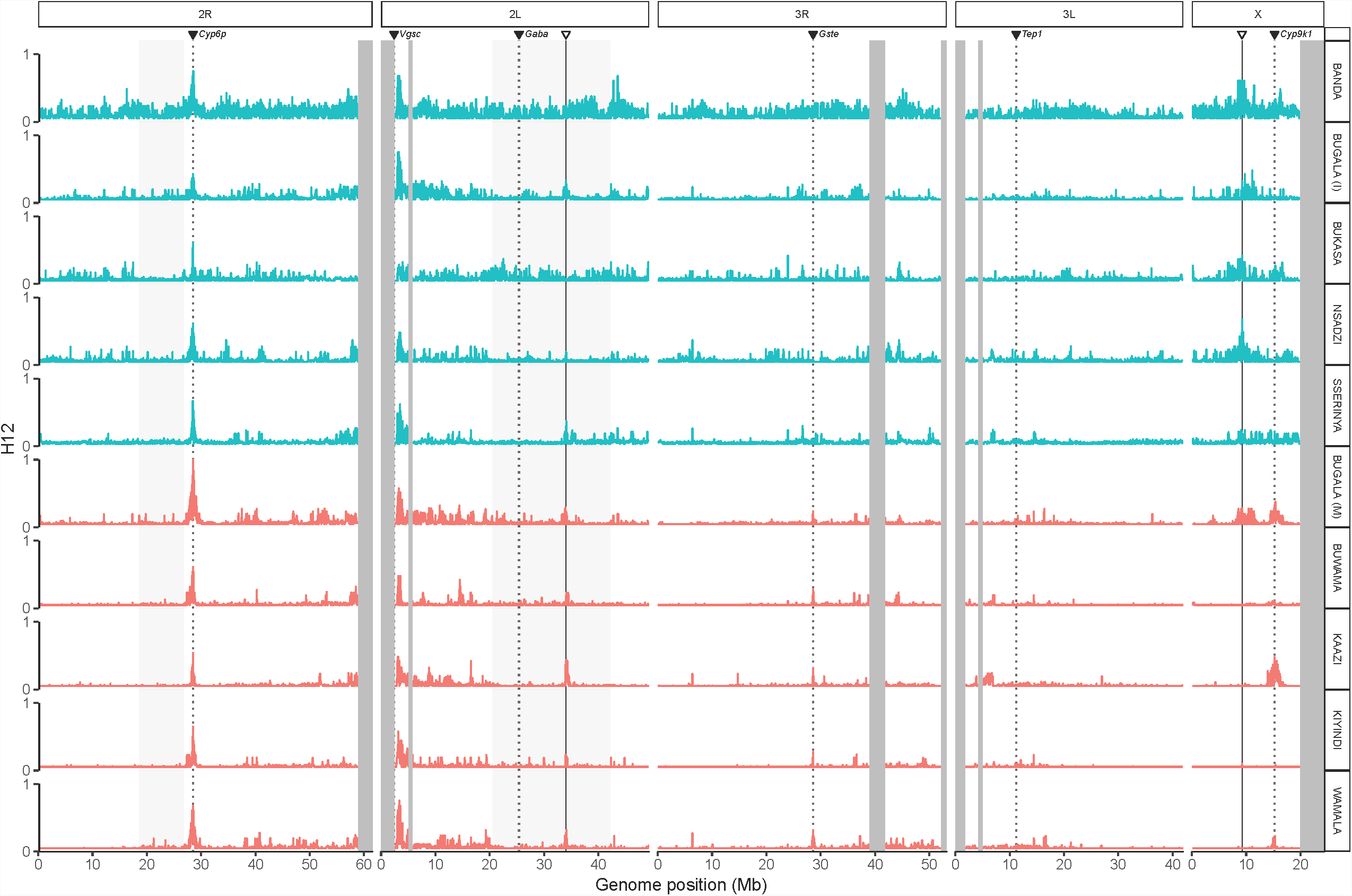
H12 across genome. Values of H12, a measure of haplotype homozygosity, plotted across genome. Shaded regions indicate inversions or heterochromatic regions (excluded from analysis) and dotted lines indicate known insecticide genes while solid lines indicate the two putative sweeps identified in the present study.

**Figure S10:**
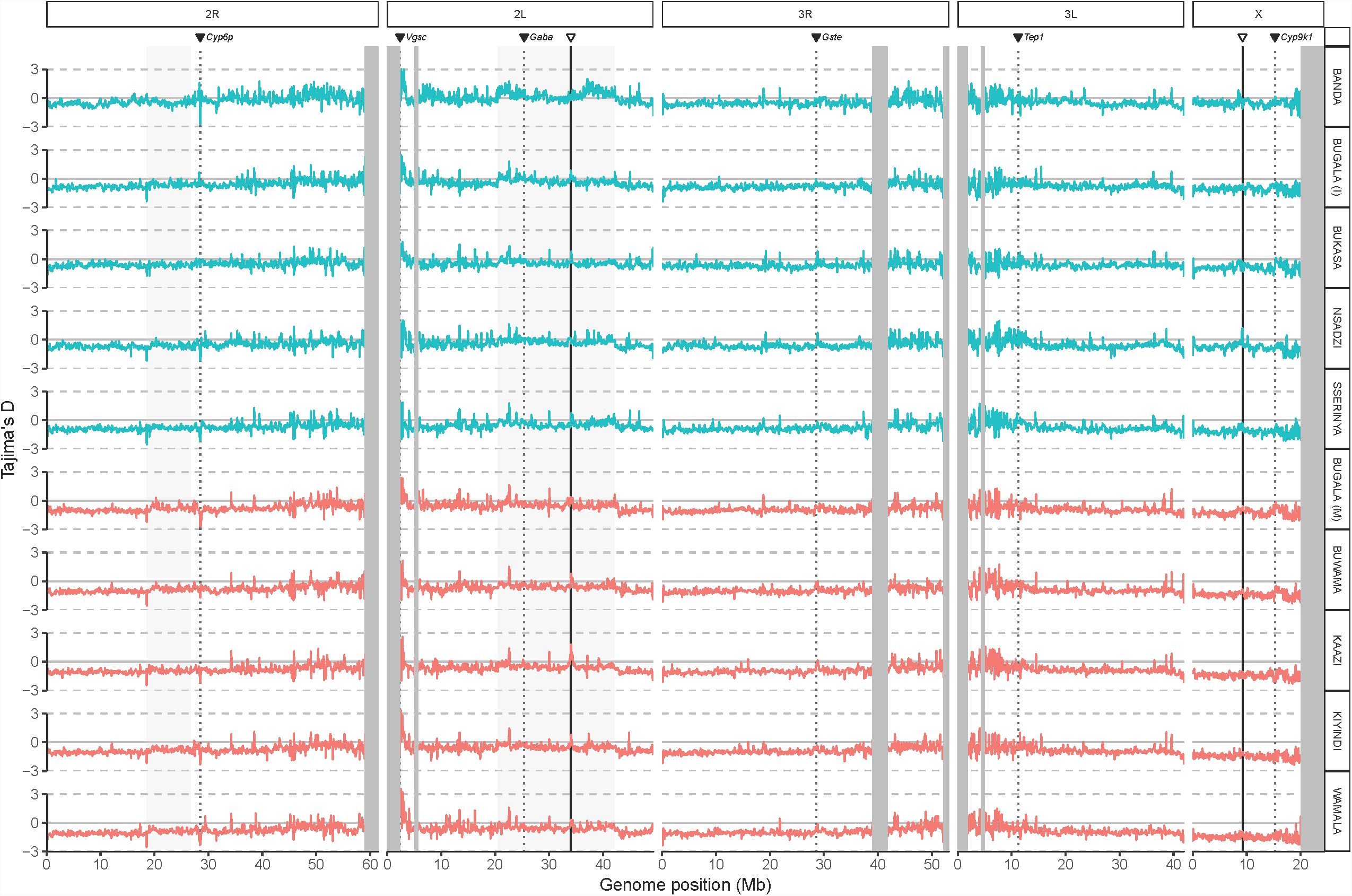
Tajima’s *D* across genome. Tajima’s *D* plotted across genome. Shaded regions indicate inversions or heterochromatic regions (excluded from analysis) and dotted lines indicate known insecticide genes while solid lines indicate the two putative sweeps identified in the present study.

**Figure S11:**
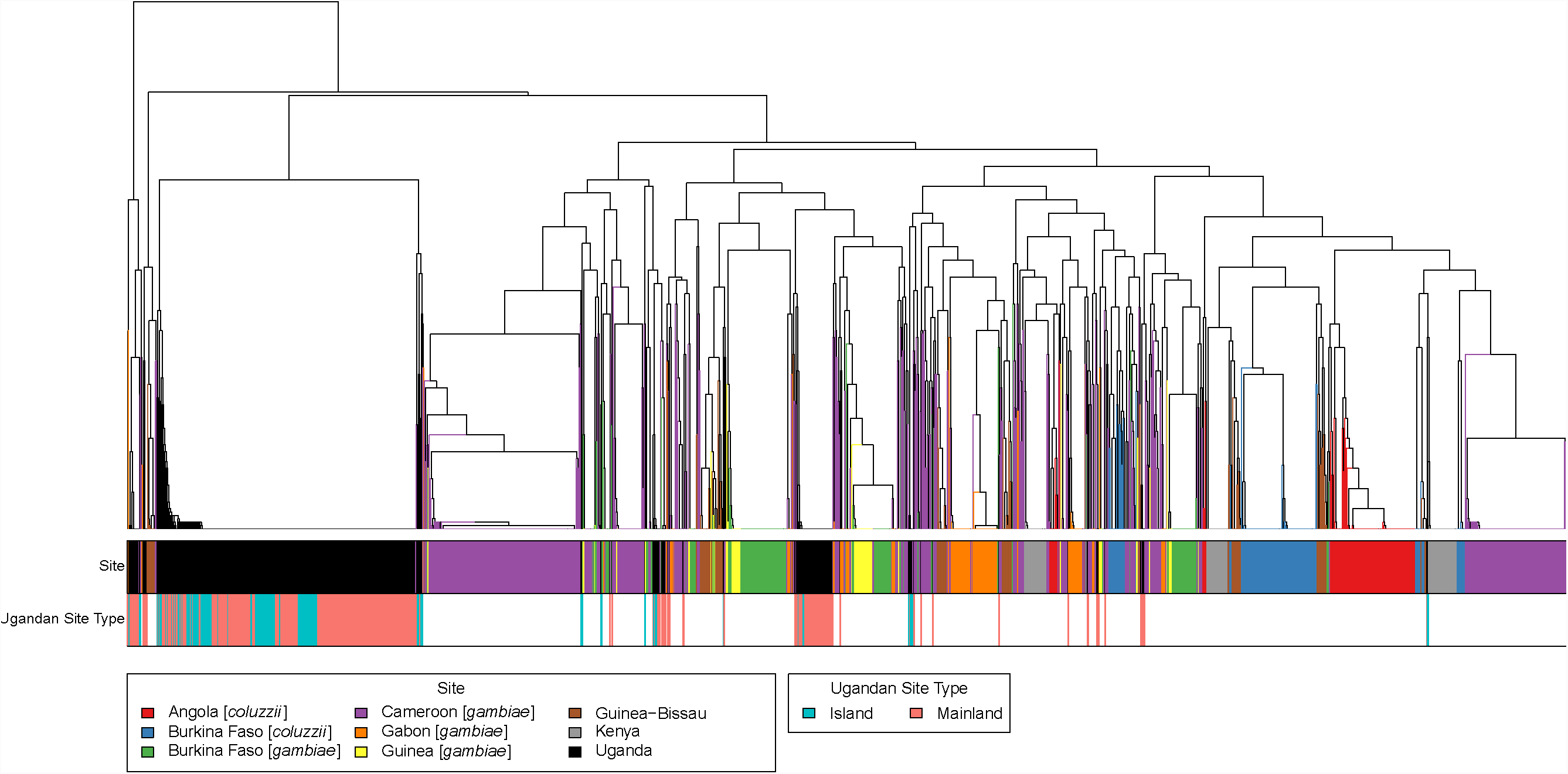
Tree for sweep on *Cyp6p* gene cluster on chromosome arm 2R. Distance-based tree of haplotypes near sweep at *Cyp6p* gene cluster on chromosome arm 2R. Region shown is 10 kb up- and downstream of sweep target, centered at chr2R:28,501,972 (the approximate location of the peaks in pairwise statistics). Top color bar indicates locality, with all Ugandan individuals, from both the Ag1000G reference population and the LVB, in black. The bottom color bar differentiates the Ugandan individuals into mainland (red) and island (blue) individuals.

**Figure S12:**
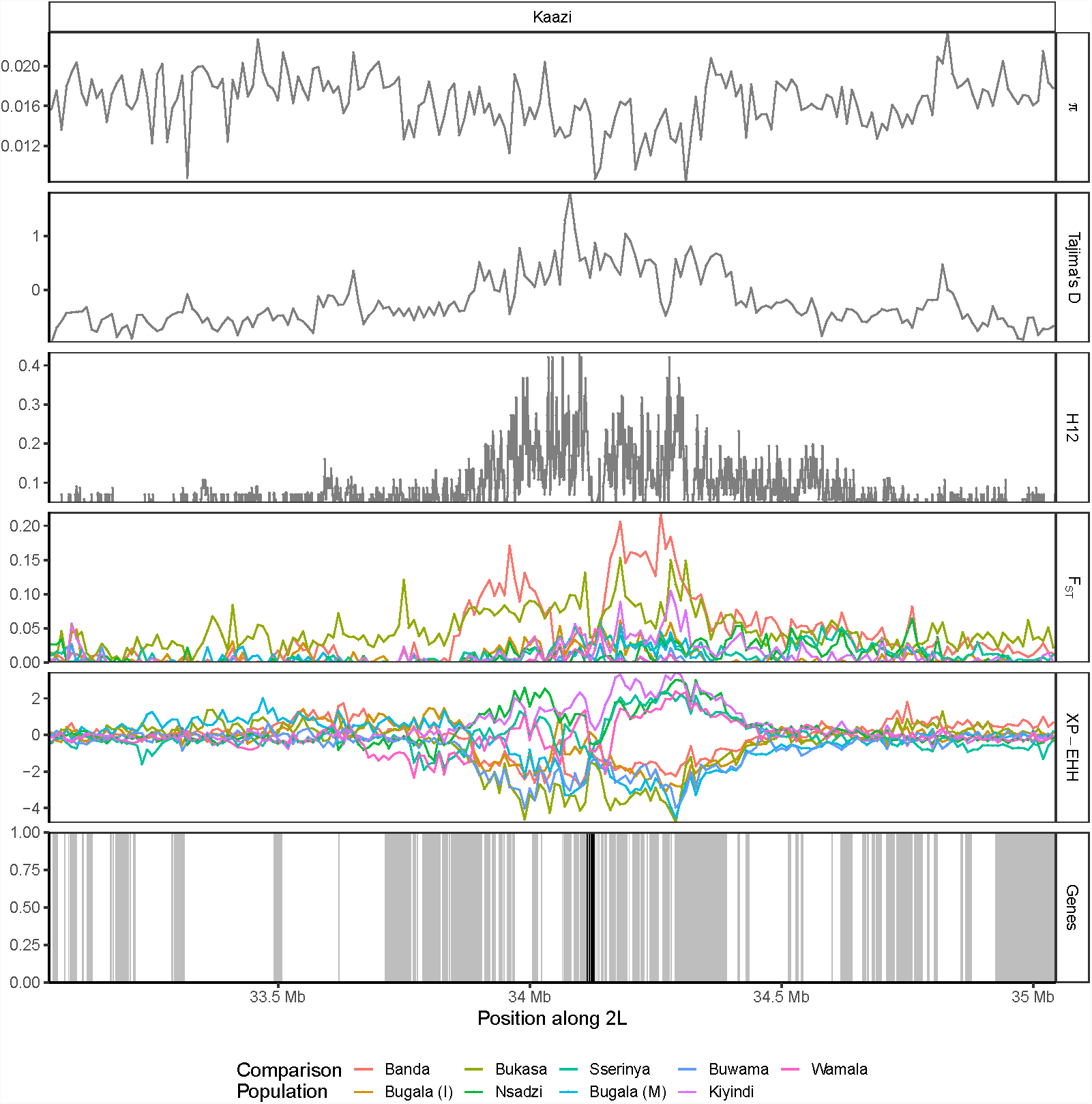
Selective sweep signal on chromosome 2L. Population genetic statistics plotted near putative sweep on chromosome 2L. Focus population for all pairwise *F*_*ST*_ and XP-EHH comparisons is mainland site Kaazi, chosen to maximize peak height in these statistics. Region shown is 1 Mb up- and downstream of sweep target, centered at chr2L:34,044,820. Several genes involved in chorion formation (AGAP006549, AGAP006550, AGAP006551, AGAP006553, AGAP006554, AGAP006555 and AGAP006556) are highlighted with black vertical lines, while other genes are indicated with gray vertical lines.

**Figure S13:**
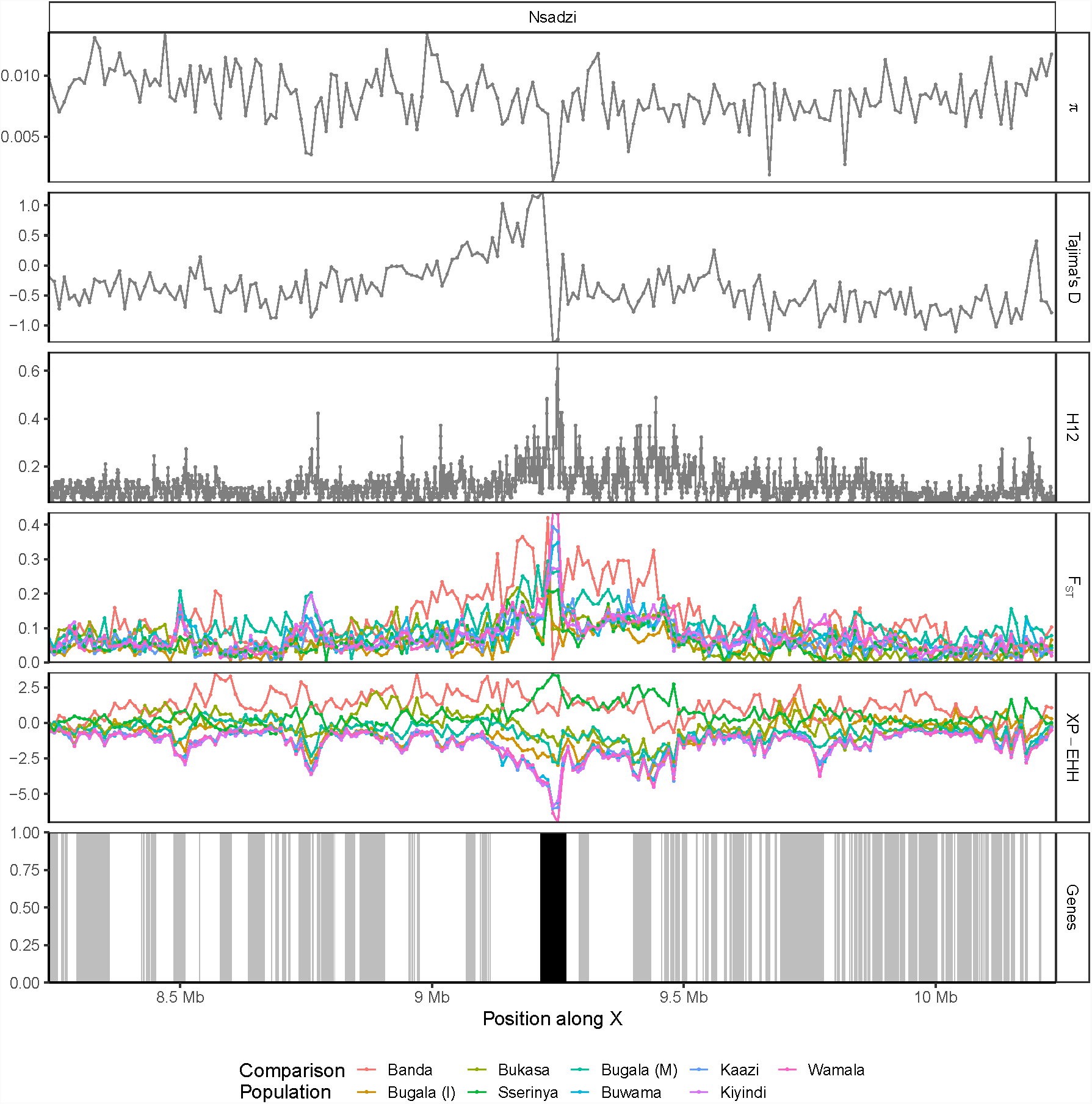
Selective sweep signal on X-chromosome near *rdgA* ortholog. Population genetic statistics plotted near putative sweep on X-chromosome. Focus population for all pairwise *F*_*ST*_ and XP-EHH comparisons is island site Nsadzi, chosen to maximize peak height in these statistics. Region shown is 1 Mb up- and downstream of sweep target, centered at chrX:9,238,942 (the approximate peak in pairwise statistics). The gene eye-specific diacylglycerol kinase (AGAP000519, chrX:9,215,505-9,266,532) is highlighted with a black vertical line, while other genes are indicated with gray vertical lines.

**Figure S14:**
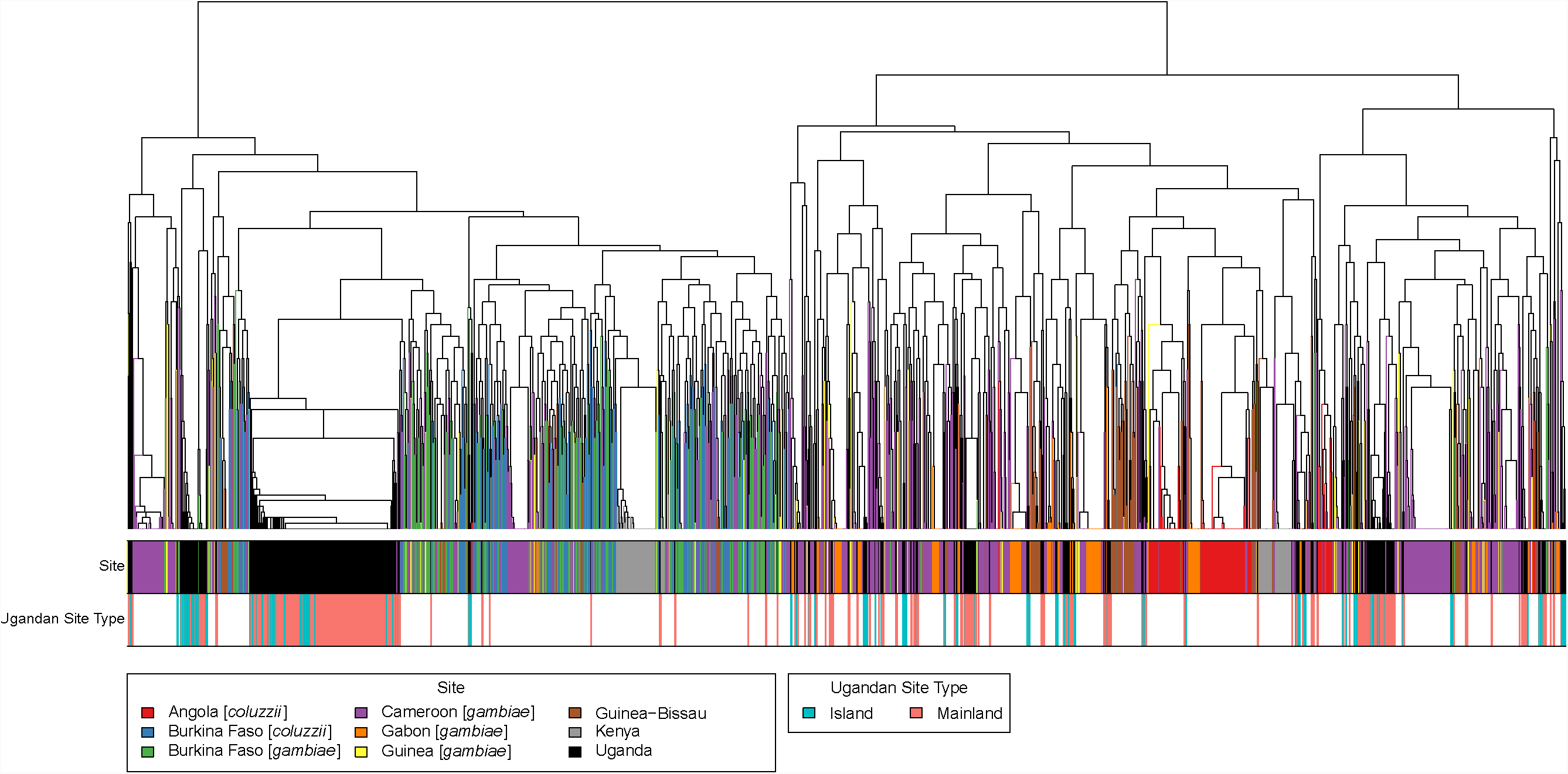
Tree for putative sweep on chromosome 2L. Distance-based tree of haplotypes near putative sweep on chromosome 2L. Region shown is 10 kb up- and downstream of sweep target, centered at chr2L:34,044,820 (the approximate location of the peaks in pairwise statistics). Top color bar indicates locality, with all Ugandan individuals, from both the Ag1000G reference population and the LVB, in black. The bottom color bar differentiates the Ugandan individuals into mainland (red) and island (blue) individuals.

**Figure S15:**
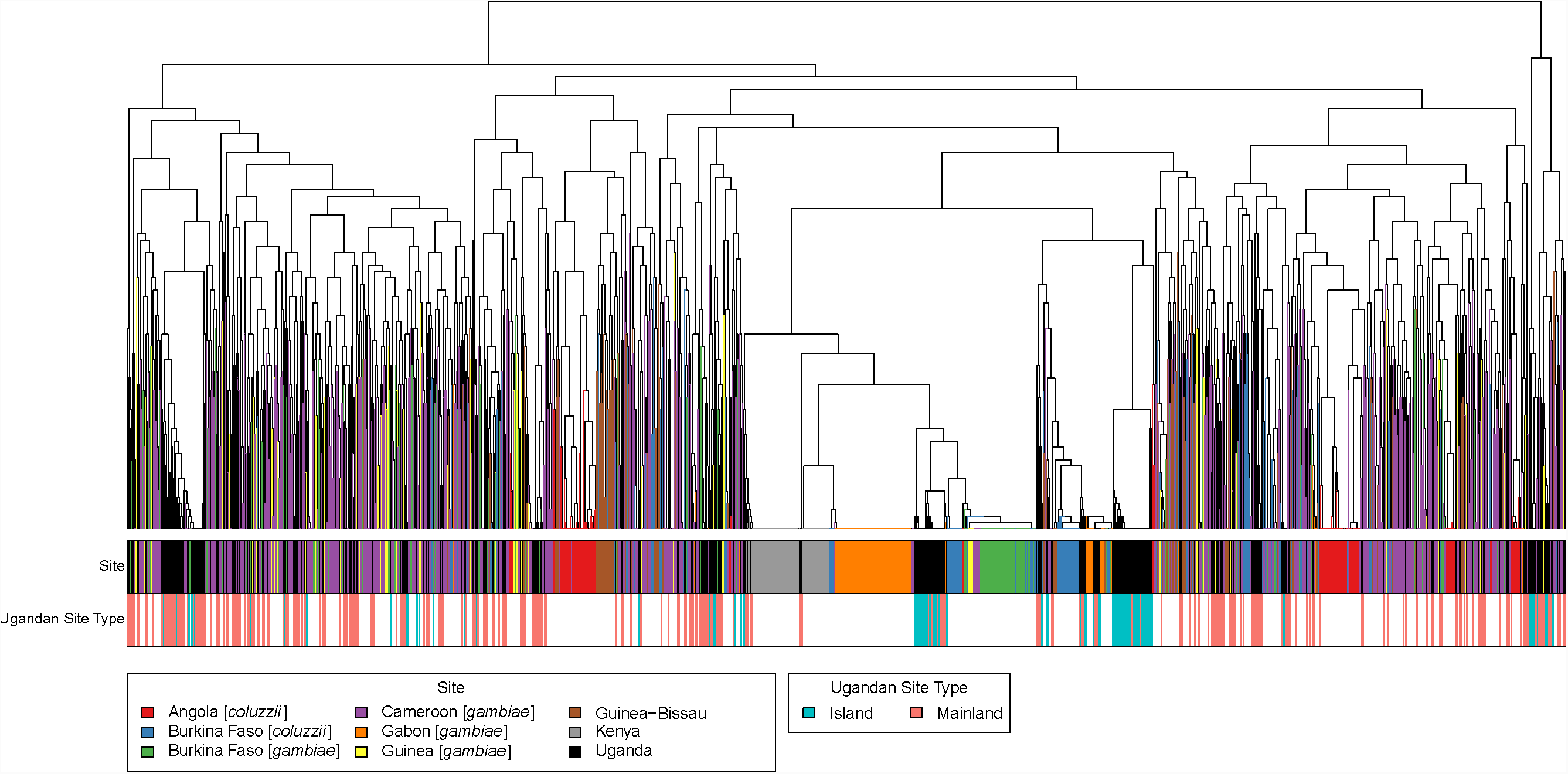
Tree for putative sweep on X-chromosome near *rdgA* ortholog. Distance-based tree of haplotypes near putative sweep on X-chromosome. Region shown is 10 kb up- and downstream of sweep target, centered at chrX:9,238,942 (the approximate location of the peaks in pairwise statistics). Top color bar indicates locality, with all Ugandan individuals, from both the Ag1000G reference population and the LVB, in black. The bottom color bar differentiates the Ugandan individuals into mainland (red) and island (blue) individuals.

**Table S1:** Sampling sites and coordinates.

| Location | Latitude | Longitude | Sample Count |
| --- | --- | --- | --- |
| Banda | -0.25893 | 32.39594 | 11 |
| Bugala - Bugoma | -0.26697 | 32.07936 | 11 |
| Bugala - Lutoboka | -0.31624 | 32.29246 | 7 |
| Bugala - Mweena | -0.32806 | 32.31113 | 5 |
| Bukasa | -0.48609 | 32.45091 | 11 |
| Buwama | 0.02077 | 32.10574 | 11 |
| Kaazi | -0.31831 | 31.88183 | 11 |
| Kiyindi | 0.27558 | 33.14699 | 10 |
| Nsadzi | -0.08632 | 32.58895 | 11 |
| Sserinya | -0.26476 | 32.37228 | 16 |
| Wamala | 0.40811 | 31.99609 | 11 |

**Table S2:** List of individuals included in study with mean depth of sequencing coverage.

| ID | Field ID | Island | Site | Mean depth |
| --- | --- | --- | --- | --- |
| LVB2015-1 | CM-KSB-J5 | Nsadzi | Kansambwe | 20.40 |
| LVB2015-2 | K-KSB-E1 | Nsadzi | Kansambwe | 24.50 |
| LVB2015-3 | RM-KSB-G1 | Nsadzi | Kansambwe | 17.90 |
| LVB2015-4 | NKG-F-G3 | Bukasa | Nakibanga | 4.90 |
| LVB2015-6 | NKG-F-H1 | Bukasa | Nakibanga | 15.80 |
| LVB2015-7 | NKG-K-I1 | Bukasa | Nakibanga | 15.80 |
| LVB2015-8 | NKG-K-K1 | Bukasa | Nakibanga | 19.90 |
| LVB2015-9 | NKG-M-C1 | Bukasa | Nakibanga | 22.30 |
| LVB2015-10 | NKG-M-D1 | Bukasa | Nakibanga | 20.30 |
| LVB2015-11 | NKG-M-F1 | Bukasa | Nakibanga | 23.90 |
| LVB2015-14 | MWN-K-A1 | Bugala | Mweena | 21.20 |
| LVB2015-15 | MWN-K-C2 | Bugala | Mweena | 18.30 |
| LVB2015-16 | MWN-P-D1 | Bugala | Mweena | 5.34 |
| LVB2015-17 | MWN-R-E1 | Bugala | Mweena | 22.60 |
| LVB2015-18 | MWN-R-F1 | Bugala | Mweena | 17.10 |
| LVB2015-19 | BDA-K-B1 | Banda | Banda | 14.40 |
| LVB2015-20 | BDA-K-B2 | Banda | Banda | 18.00 |
| LVB2015-21 | BBS-C-M1 | Sserinya | Bbosa | 19.70 |
| LVB2015-22 | BBS-F-F1 | Sserinya | Bbosa | 20.80 |
| LVB2015-24 | BBS-K-J3 | Sserinya | Bbosa | 22.80 |
| LVB2015-25 | BBS-K-J8 | Sserinya | Bbosa | 17.30 |
| LVB2015-26 | BBS-K-K2 | Sserinya | Bbosa | 18.20 |
| LVB2015-27 | BBS-M-L1 | Sserinya | Bbosa | 22.40 |
| LVB2015-28 | BBS-P-I4 | Sserinya | Bbosa | 23.30 |
| LVB2015-29 | BBS-R-A2 | Sserinya | Bbosa | 19.60 |
| LVB2015-30 | BBS-R-C1 | Sserinya | Bbosa | 16.10 |
| LVB2015-32 | KSS-F-E2 | Sserinya | Kasisa | 21.00 |
| LVB2015-33 | LBK-C-F1 | Bugala | Lutoboka | 21.00 |
| LVB2015-34 | LBK-C-F6 | Bugala | Lutoboka | 18.60 |
| LVB2015-35 | LBK-C-G6 | Bugala | Lutoboka | 20.90 |
| LVB2015-36 | LBK-K-E2 | Bugala | Lutoboka | 22.00 |
| LVB2015-37 | LBK-M-A1 | Bugala | Lutoboka | 18.50 |
| LVB2015-39 | LBK-R-O1 | Bugala | Lutoboka | 20.20 |
| LVB2015-42 | BGM-F-D1 | Bugala | Bugoma | 16.70 |
| LVB2015-43 | BGM-F-E2 | Bugala | Bugoma | 24.50 |
| LVB2015-45 | BGM-K-M2 | Bugala | Bugoma | 23.80 |
| LVB2015-46 | BGM-M-G1 | Bugala | Bugoma | 18.40 |
| LVB2015-47 | BGM-M-H2 | Bugala | Bugoma | 14.10 |
| LVB2015-48 | BGM-M-J1 | Bugala | Bugoma | 20.90 |
| LVB2015-50 | BGM-P-F9 | Bugala | Bugoma | 18.90 |
| LVB2015-51 | BGM-R-O2 | Bugala | Bugoma | 16.80 |
| LVB2015-52 | KZI-F-F001 | Kaazi | Nabugabo | 19.10 |
| LVB2015-53 | KZI-F-G001 | Kaazi | Nabugabo | 19.50 |
| LVB2015-54 | KZI-F-H001 | Kaazi | Nabugabo | 19.40 |
| LVB2015-55 | KZI-P-A001 | Kaazi | Nabugabo | 10.30 |
| LVB2015-56 | KZI-P-B005 | Kaazi | Nabugabo | 16.50 |
| LVB2015-59 | KZI-R-C003 | Kaazi | Nabugabo | 18.10 |
| LVB2015-60 | KZI-R-D007 | Kaazi | Nabugabo | 15.90 |
| LVB2015-61 | BWM-C-G001 | Buwama | Buwama | 16.10 |
| LVB2015-62 | BWM-C-H001 | Buwama | Buwama | 11.10 |
| LVB2015-63 | BWM-F-A001 | Buwama | Buwama | 20.40 |
| LVB2015-64 | BWM-F-B001 | Buwama | Buwama | 21.50 |
| LVB2015-65 | BWM-P-J001 | Buwama | Buwama | 14.80 |
| LVB2015-66 | BWM-R-C002 | Buwama | Buwama | 19.10 |
| LVB2015-67 | BWM-R-F005 | Buwama | Buwama | 22.30 |
| LVB2015-68 | NMA-C-E003 | Wamala | Naama | 20.30 |
| LVB2015-69 | NMA-C-F002 | Wamala | Naama | 17.80 |
| LVB2015-70 | NMA-F-A001 | Wamala | Naama | 13.10 |
| LVB2015-71 | NMA-K-B001 | Wamala | Naama | 22.10 |
| LVB2015-72 | NMA-K-C002 | Wamala | Naama | 18.00 |
| LVB2015-73 | NMA-P-G001 | Wamala | Naama | 18.20 |
| LVB2015-74 | NMA-P-H003 | Wamala | Naama | 16.60 |
| LVB2015-76 | KYD-C-G001 | Kiyindi | Kiyindi | 16.10 |
| LVB2015-77 | KYD-C-H001 | Kiyindi | Kiyindi | 11.80 |
| LVB2015-78 | KYD-C-I001 | Kiyindi | Kiyindi | 16.40 |
| LVB2015-79 | KYD-C-J002 | Kiyindi | Kiyindi | 11.50 |
| LVB2015-80 | KYD-F-A003 | Kiyindi | Kiyindi | 10.30 |
| LVB2015-81 | KYD-F-B004 | Kiyindi | Kiyindi | 21.50 |
| LVB2015-82 | KYD-K-D002 | Kiyindi | Kiyindi | 18.40 |
| LVB2015-84 | KYD-R-K001 | Kiyindi | Kiyindi | 16.80 |
| LVB2015-89 | BDA-K-E2 | Banda | Banda | 15.10 |
| LVB2015-90 | BDA-K-F1 | Banda | Banda | 25.10 |
| LVB2015-91 | BDA-M-N1 | Banda | Banda | 25.60 |
| LVB2015-92 | BDA-M-O4 | Banda | Banda | 17.60 |
| LVB2015-93 | BDA-M-Q1 | Banda | Banda | 39.20 |
| LVB2015-96 | CM-KSB-J2 | Nsadzi | Kansambwe | 9.22 |
| LVB2015-97 | CM-KSB-J3 | Nsadzi | Kansambwe | 10.10 |
| LVB2015-98 | CM-KSB-J6 | Nsadzi | Kansambwe | 16.90 |
| LVB2015-100 | K-KSB-D1 | Nsadzi | Kansambwe | 6.05 |
| LVB2015-101 | ML-KSB-M1 | Nsadzi | Kansambwe | 4.27 |
| LVB2015-102 | ML-KSB-M2 | Nsadzi | Kansambwe | 19.90 |
| LVB2015-103 | RM-KSB-G2 | Nsadzi | Kansambwe | 14.20 |
| LVB2015-104 | RM-KSB-G3 | Nsadzi | Kansambwe | 17.50 |
| LVB2015-105 | NKG-R-A12 | Bukasa | Nakibanga | 15.30 |
| LVB2015-106 | NKG-C-E1 | Bukasa | Nakibanga | 16.20 |
| LVB2015-108 | NKG-K-C5 | Bukasa | Nakibanga | 18.50 |
| LVB2015-109 | NKG-M-A1 | Bukasa | Nakibanga | 12.80 |
| LVB2015-112 | BDA-K-D4 | Banda | Banda | 12.70 |
| LVB2015-113 | BDA-K-E3 | Banda | Banda | 12.20 |
| LVB2015-114 | BDA-M-N5 | Banda | Banda | 15.00 |
| LVB2015-115 | BDA-M-P1 | Banda | Banda | 16.80 |
| LVB2015-116 | BBS-C-M3 | Sserinya | Bbosa | 16.60 |
| LVB2015-117 | BBS-K-J1 | Sserinya | Bbosa | 18.80 |
| LVB2015-118 | BBS-K-J11 | Sserinya | Bbosa | 14.60 |
| LVB2015-120 | BBS-K-K6 | Sserinya | Bbosa | 18.10 |
| LVB2015-121 | BBS-P-I8 | Sserinya | Bbosa | 15.00 |
| LVB2015-122 | BBS-R-A19 | Sserinya | Bbosa | 15.50 |
| LVB2015-125 | LBK-R-A5 | Bugala | Lutoboka | 18.10 |
| LVB2015-126 | BGM-K-K1 | Bugala | Bugoma | 15.20 |
| LVB2015-128 | BGM-M-H4 | Bugala | Bugoma | 20.30 |
| LVB2015-129 | BGM-P-F4 | Bugala | Bugoma | 19.00 |
| LVB2015-130 | KZI-F-G005 | Kaazi | Nabugabo | 18.60 |
| LVB2015-131 | KZI-P-A007 | Kaazi | Nabugabo | 15.80 |
| LVB2015-132 | KZI-R-C012 | Kaazi | Nabugabo | 15.10 |
| LVB2015-133 | KZI-R-E011 | Kaazi | Nabugabo | 16.30 |
| LVB2015-134 | BWM-P-I001 | Buwama | Buwama | 18.20 |
| LVB2015-135 | BWM-P-K002 | Buwama | Buwama | 19.30 |
| LVB2015-136 | BWM-R-D001 | Buwama | Buwama | 14.40 |
| LVB2015-137 | BWM-R-F002 | Buwama | Buwama | 19.90 |
| LVB2015-138 | NMA-C-E006 | Wamala | Naama | 21.90 |
| LVB2015-139 | NMA-C-F003 | Wamala | Naama | 20.40 |
| LVB2015-140 | NMA-P-G003 | Wamala | Naama | 18.90 |
| LVB2015-141 | NMA-R-I001 | Wamala | Naama | 14.10 |
| LVB2015-142 | KYD-F-B006 | Kiyindi | Kiyindi | 18.10 |
| LVB2015-143 | KYD-K-E003 | Kiyindi | Kiyindi | 14.20 |

**Table S3:** Results of two population demographic inference with IM model in *δ*a*δ*i when comparing island to island localities. Numbers in parentheses are bounds of 95% confidence interval computed using Fisher information matrix and 100 bootstrap replicates of 1 Mb from the dataset.

| Localities | $N_a$ | % Pop. 1 in Split | Pop. 1 $\nu_F$ | Pop. 2 $\nu_F$ | Time since split | $m_{12}$ | $m_{21}$ |
| --- | --- | --- | --- | --- | --- | --- | --- |
| Banda - Bugala (I) | 531,000<br>(530,000, 532,000) | 0.603<br>(0.596, 0.609) | 1.94<br>(1.8, 2.09) | 9,800<br>(7,060, 12,500) | 3,290<br>(3,190, 3,390) | None | None |
| Banda - Bukasa | 526,000<br>(525,000, 527,000) | 0.568<br>(0.556, 0.581) | 14.7<br>(13.7, 15.6) | 9,080<br>(7,240, 10,900) | 7,580<br>(7,470, 7,690) | None | None |
| Banda - Nsadzi | 527,000<br>(526,000, 528,000) | 0.518<br>(0.502, 0.534) | 47.1<br>(39.9, 54.4) | 9,880<br>(7,310, 12,400) | 9,340<br>(9,160, 9,530) | None | None |
| Banda - Sserinya | 531,000<br>(530,000, 532,000) | 0.489<br>(0.464, 0.514) | 10.5<br>(8.69, 12.4) | 9,840<br>(7,390, 12,300) | 4,430<br>(4,250, 4,610) | None | None |
| Bugala (I) - Bukasa | 527,000<br>(526,000, 528,000) | 0.49<br>(0.471, 0.509) | 9,840<br>(7,850, 11,800) | 536<br>(447, 624) | 5,290<br>(5,170, 5,410) | None | None |
| Bugala (I) - Nsadzi | 526,000<br>(525,000, 527,000) | 0.56<br>(0.536, 0.583) | 8,980<br>(6,920, 11,000) | 128<br>(110, 146) | 6,680<br>(6,440, 6,910) | None | None |
| Bugala (I) - Sserinya | 530,000<br>(529,000, 531,000) | 0.61<br>(0.571, 0.65) | 5,850<br>(3,610, 8,090) | 49<br>(38.3, 59.6) | 2,130<br>(1,950, 2,300) | None | None |
| Bukasa - Nsadzi | 527,000<br>(526,000, 528,000) | 0.499<br>(0.49, 0.507) | 3,420<br>(2,860, 3,980) | 335<br>(293, 377) | 9,340<br>(9,190, 9,500) | None | None |
| Bukasa - Sserinya | 525,000<br>(524,000, 526,000) | 0.503<br>(0.495, 0.51) | 9,840<br>(7,490, 12,200) | 6,760<br>(5,630, 7,880) | 8,930<br>(8,780, 9,080) | None | None |
| Nsadzi - Sserinya | 540,000<br>(539,000, 541,000) | 0.538<br>(0.521, 0.554) | 893<br>(612, 1,170) | 9,900<br>(7,810, 12,000) | 9,540<br>(9,280, 9,790) | None | None |

**Table S4:** Results of two population demographic inference with IM model in *δ*a*δ*i when comparing island to mainland localities. Numbers in parentheses are bounds of 95% confidence interval computed using Fisher information matrix and 100 bootstrap replicates of 1 Mb from the dataset.

| Localities | $N_a$ | % Pop. 1 in Split | Pop. 1 $\nu_F$ | Pop. 2 $\nu_F$ | Time since split | $m_{12}$ | $m_{21}$ |
| --- | --- | --- | --- | --- | --- | --- | --- |
| Banda - Bugala (M) | 522,000<br>(521,000, 523,000) | 0.599<br>(0.586, 0.612) | 3.83<br>(3.59, 4.07) | 9,900<br>(8,410, 11,400) | 5,400<br>(5,300, 5,500) | None | None |
| Banda - Buwama | 522,000<br>(521,000, 523,000) | 0.51<br>(0.491, 0.529) | 1.09<br>(1.01, 1.16) | 8,520<br>(6,830, 10,200) | 3,040<br>(2,940, 3,150) | None | None |
| Banda - Kaazi | 522,000<br>(521,000, 523,000) | 0.563<br>(0.55, 0.575) | 3.99<br>(3.71, 4.28) | 9,960<br>(8,370, 11,500) | 5,890<br>(5,760, 6,030) | None | None |
| Banda - Kiyindi | 510,000<br>(509,000, 511,000) | 0.568<br>(0.56, 0.577) | 1.72<br>(1.69, 1.75) | 9,910<br>(8,170, 11,600) | 3,910<br>(3,880, 3,950) | None | None |
| Banda - Wamala | 521,000<br>(520,000, 522,000) | 0.562<br>(0.554, 0.57) | 3.9<br>(3.66, 4.15) | 7,970<br>(6,790, 9,140) | 5,510<br>(5,390, 5,620) | None | None |
| Bugala (I) - Bugala (M) | 522,000<br>(521,000, 523,000) | 0.555<br>(0.543, 0.566) | 1,440<br>(954, 1,930) | 9,710<br>(7,420, 12,000) | 4,170<br>(4,040, 4,300) | None | None |
| Bugala (I) - Buwama | 523,000<br>(522,000, 524,000) | 0.499<br>(0.496, 0.503) | 0.215<br>(0.209, 0.222) | 130<br>(2.12, 258) | 190<br>(188, 192) | None | None |
| Bugala (I) - Kaazi | 523,000<br>(522,000, 524,000) | 0.366<br>(0.348, 0.384) | 1,420<br>(1,210, 1,620) | 8,510<br>(6,610, 10,400) | 5,060<br>(4,890, 5,230) | None | None |
| Bugala (I) - Kiyindi | 508,000<br>(507,000, 509,000) | 0.363<br>(0.342, 0.383) | 1,350<br>(1,070, 1,620) | 5,910<br>(4,470, 7,350) | 3,580<br>(3,420, 3,740) | None | None |
| Bugala (I) - Wamala | 520,000<br>(519,000, 521,000) | 0.486<br>(0.462, 0.51) | 2,060<br>(1,610, 2,500) | 9,270<br>(6,790, 11,800) | 3,700<br>(3,570, 3,830) | None | None |
| Bugala (M) - Bukasa | 521,000<br>(520,000, 522,000) | 0.53<br>(0.508, 0.552) | 9,000<br>(6,720, 11,300) | 66.3<br>(52.2, 80.5) | 4,830<br>(4,670, 4,990) | None | None |
| Bugala (M) - Nsadzi | 535,000 | 0.439 | 9,220 | 7.59 | 4,500 | None | None |
|  | (534,000, 536,000) | (0.423, 0.455) | (7,500, 10,900) | (6.91, 8.26) | (4,390, 4,620) |  |  |
| Bugala (M) - Sserinya | 534,000 | 0.516 | 9,590 | 48.8 | 4,040 | None | None |
|  | (533,000, 535,000) | (0.497, 0.535) | (7,640, 11,500) | (42.4, 55.2) | (3,930, 4,160) |  |  |
| Bukasa - Buwama | 522,000 | 0.501 | 1.43 | 9,960 | 1,690 | None | None |
|  | (521,000, 523,000) | (0.496, 0.506) | (1.4, 1.46) | (7,250, 12,700) | (1,670, 1,710) |  |  |
| Bukasa - Kaazi | 522,000 | 0.501 | 41.5 | 9,410 | 5,490 | None | None |
|  | (521,000, 522,000) | (0.491, 0.51) | (33.8, 49.3) | (6,990, 11,800) | (5,320, 5,650) |  |  |
| Bukasa - Kiyindi | 508,000 | 0.361 | 18.6 | 9,220 | 3,360 | None | None |
|  | (507,000, 509,000) | (0.336, 0.386) | (15.8, 21.4) | (6,790, 11,600) | (3,190, 3,540) |  |  |
| Bukasa - Wamala | 520,000 | 0.39 | 304 | 8,060 | 5,200 | None | None |
|  | (519,000, 521,000) | (0.372, 0.409) | (257, 351) | (6,270, 9,850) | (5,050, 5,360) |  |  |
| Buwama - Nsadzi | 524,000 | 0.54 | 9,810 | 1.25 | 1,930 | None | None |
|  | (523,000, 525,000) | (0.502, 0.578) | (6,950, 12,700) | (1.07, 1.44) | (1,800, 2,070) |  |  |
| Buwama - Sserinya | 523,000 | 0.493 | 54.2 | 0.137 | 187 | None | None |
|  | (522,000, 524,000) | (0.488, 0.498) | (24.2, 84.2) | (0.134, 0.14) | (186, 189) |  |  |
| Kaazi - Nsadzi | 524,000 | 0.489 | 9,540 | 19.4 | 5,730 | None | None |
|  | (523,000, 525,000) | (0.47, 0.507) | (7,690, 11,400) | (16, 22.8) | (5,540, 5,910) |  |  |
| Kaazi - Sserinya | 523,000 | 0.598 | 6,980 | 104 | 4,550 | None | None |
|  | (522,000, 524,000) | (0.579, 0.618) | (5,440, 8,510) | (89.7, 119) | (4,410, 4,680) |  |  |
| Kiyindi - Nsadzi | 511,000 | 0.499 | 9,940 | 2.62 | 2,840 | None | None |
|  | (510,000, 512,000) | (0.491, 0.507) | (7,340, 12,500) | (2.26, 2.98) | (2,690, 2,990) |  |  |
| Kiyindi - Sserinya | 509,000 | 0.5 | 1,870 | 0.129 | 185 | None | None |
|  | (508,000, 510,000) | (0.496, 0.503) | (-5,010, 8,750) | (0.126, 0.132) | (183, 187) |  |  |
| Wamala - Nsadzi | 523,000 | 0.473 | 8,490 | 9.72 | 4,420 | None | None |
|  | (522,000, 524,000) | (0.454, 0.492) | (6,720, 10,200) | (8.57, 10.9) | (4,290, 4,560) |  |  |
| Wamala - Sserinya | 521,000 | 0.649 | 9,840 | 39.1 | 2,790 | None | None |
|  | (520,000, 522,000) | (0.643, 0.655) | (7,070, 12,600) | (38.1, 40.1) | (2,760, 2,820) |  |  |

**Table S5:** Results of two population demographic inference with IM model in *δ*a*δ*i when comparing mainland to mainland localities. Numbers in parentheses are bounds of 95% confidence interval computed using Fisher information matrix and 100 bootstrap replicates of 1 Mb from the dataset.

| Localities | $N_a$ | % Pop. 1 in Split | Pop. 1 $\nu_F$ | Pop. 2 $\nu_F$ | Time since split | $m_{12}$ | $m_{21}$ |
| --- | --- | --- | --- | --- | --- | --- | --- |
| Bugala (M) - Buwama | 523,000<br>(523,000, 524,000) | 0.498<br>(0.49, 0.506) | 9,590<br>(802, 18,400) | 1,090<br>(573, 1,620) | 368<br>(320, 416) | None | None |
| Bugala (M) - Kaazi | 521,000<br>(520,000, 522,000) | 0.483<br>(0.458, 0.509) | 7,570<br>(5,740, 9,400) | 8,450<br>(5,860, 11,000) | 2,710<br>(2,590, 2,830) | None | None |
| Bugala (M) - Kiyindi | 507,000<br>(506,000, 508,000) | 0.361<br>(0.318, 0.404) | 1.88<br>(1.74, 2.01) | 3,870<br>(1,810, 5,920) | 355<br>(348, 362) | 0.0000169<br>(-14.3, 14.3) | 0<br>(-31,100, 31,100) |
| Bugala (M) - Wamala | 521,000<br>(520,000, 521,000) | 0.51<br>(0.491, 0.53) | 13.1<br>(9.03, 17.1) | 323<br>(-168, 814) | 186<br>(179, 194) | 0<br>(-162, 162) | 0.00226<br>(-3,980, 3,980) |
| Buwama - Kaazi | 523,000<br>(522,000, 524,000) | 0.553<br>(0.357, 0.749) | 706<br>(-764, 2,180) | 186<br>(-101, 472) | 181<br>(122, 239) | 3.16<br>(-5,690, 5,690) | 2.12<br>(-836, 841) |
| Buwama - Kiyindi | 511,000<br>(510,000, 512,000) | 0.322<br>(0.287, 0.357) | 9,110<br>(5,390, 12,800) | 9,810<br>(4,390, 15,200) | 369<br>(302, 436) | 761<br>(-80,200, 81,700) | 178<br>(-81,200, 81,500) |
| Buwama - Wamala | 523,000<br>(522,000, 524,000) | 0.365<br>(-1,000, 1,000) | 31.7<br>(22, 41.3) | 19.7<br>(13.6, 25.8) | 124<br>(107, 141) | 9.27<br>(-578, 596) | 0<br>(-344, 344) |
| Kaazi - Kiyindi | 510,000<br>(509,000, 511,000) | 0.448<br>(0.325, 0.571) | 9,900<br>(1,100, 18,700) | 1,670<br>(749, 2,600) | 438<br>(365, 510) | 923<br>(-64,900, 66,800) | 718<br>(610, 825) |
| Kaazi - Wamala | 521,000<br>(520,000, 522,000) | 0.581<br>(0.515, 0.647) | 9,200<br>(3,680, 14,700) | 4,820<br>(2,430, 7,210) | 974<br>(795, 1,150) | None | None |
| Kiyindi - Wamala | 510,000<br>(509,000, 511,000) | 0.454<br>(0.391, 0.518) | 134<br>(57.9, 209) | 40.3<br>(25.4, 55.3) | 181<br>(166, 197) | 0<br>(-663, 663) | 0.00363<br>(-3.69, 3.7) |

**Table S6:**
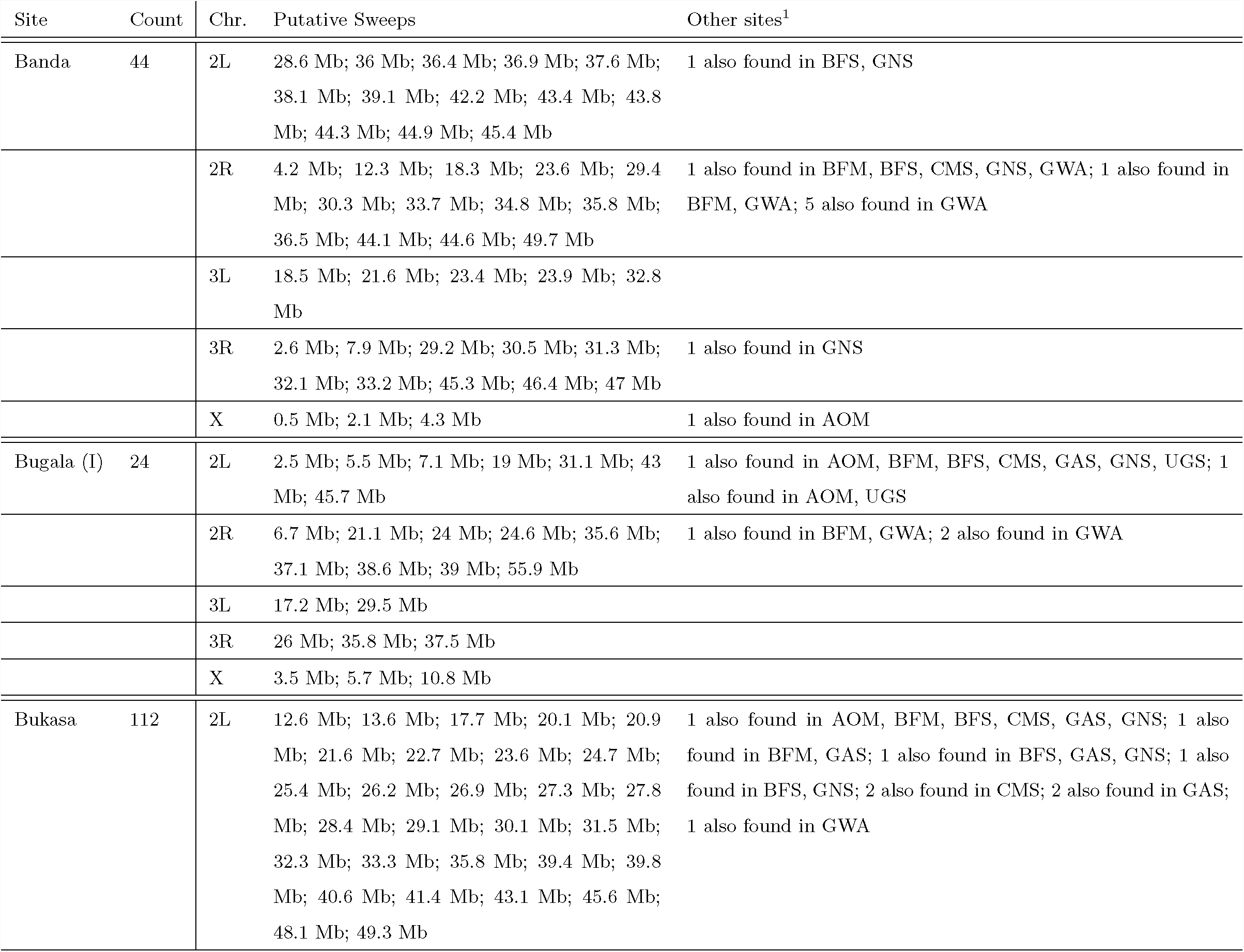
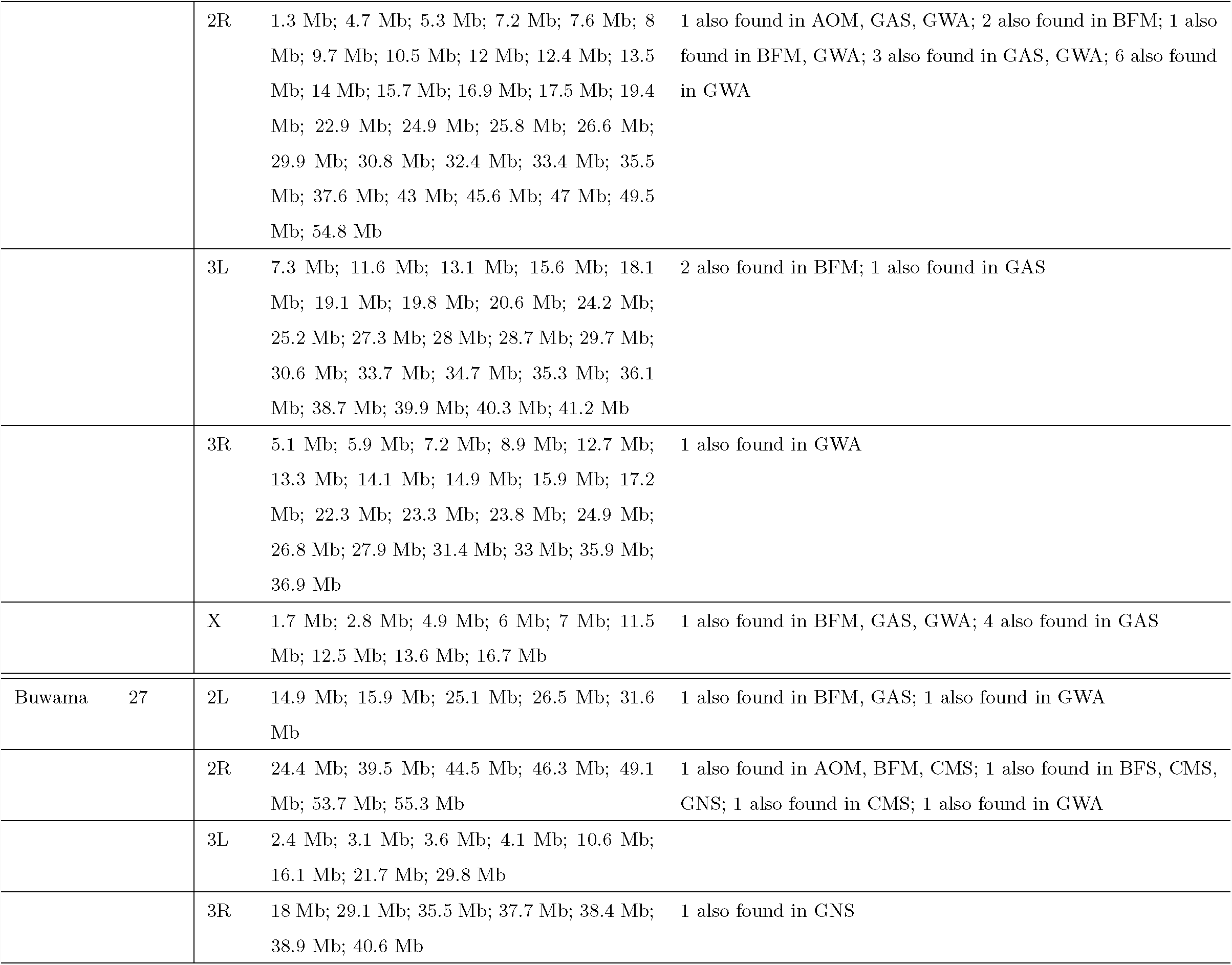
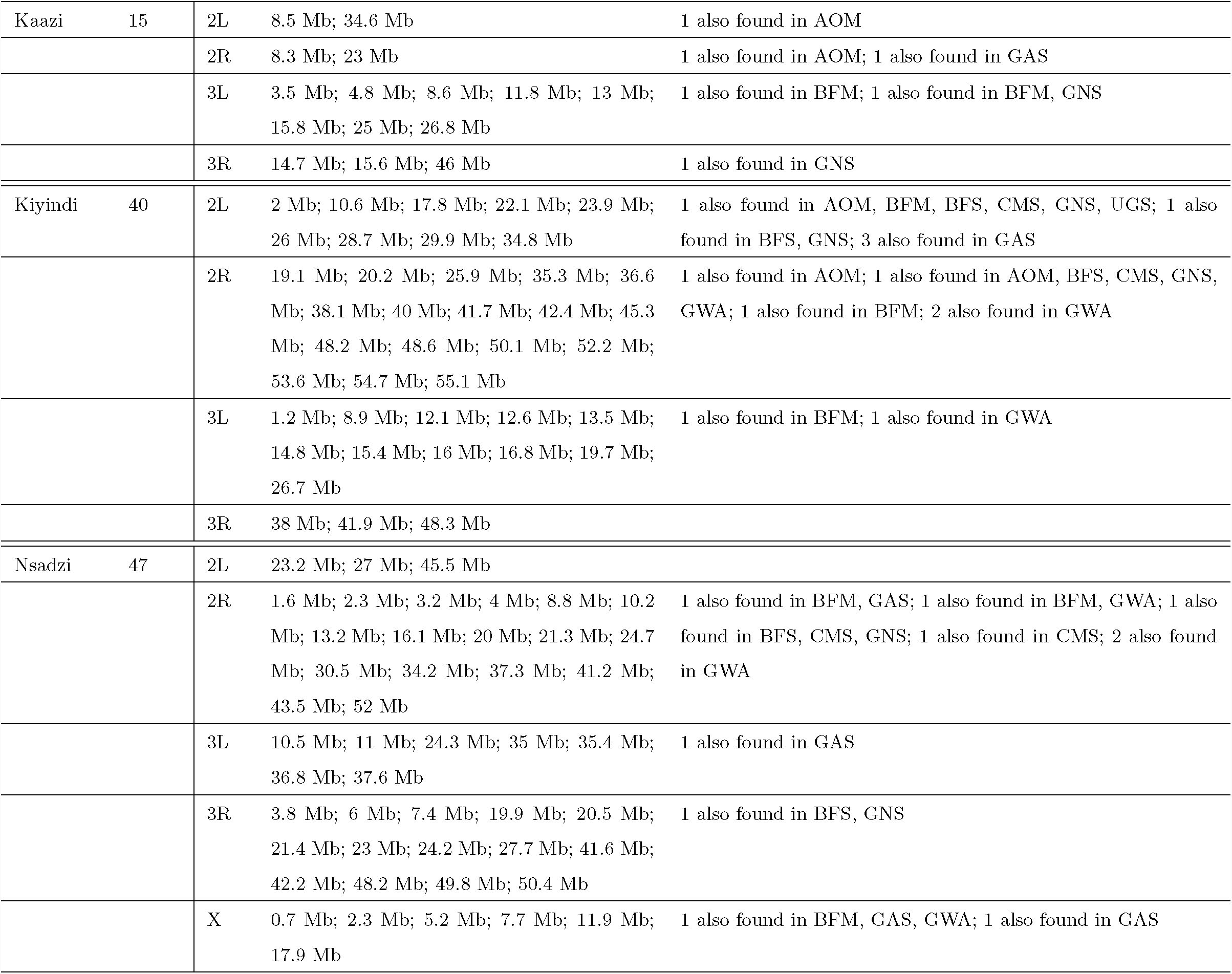
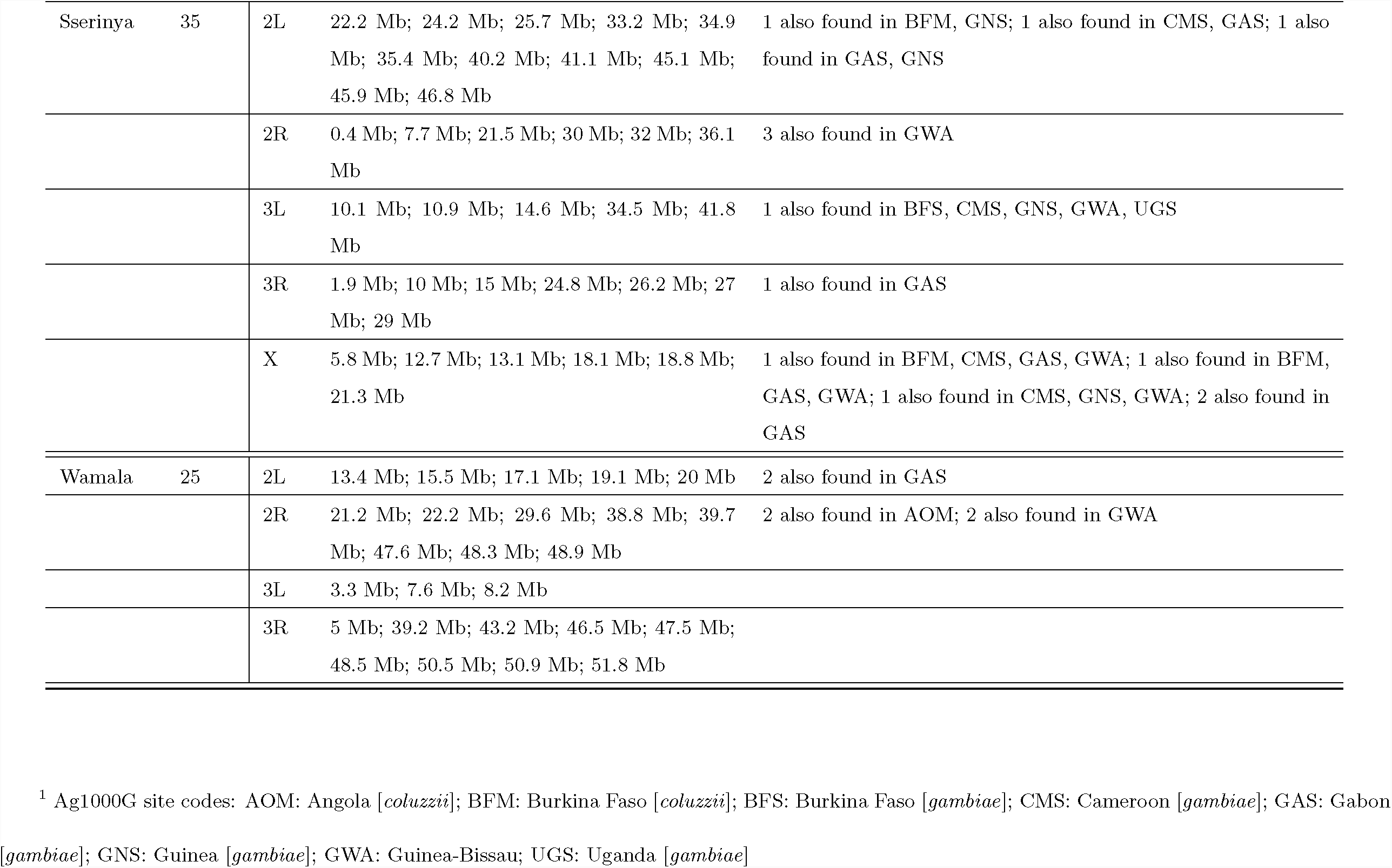
Locality-specific (in LVB) putative sweeps based on H12 statistic.

**Table S7:** Putative sweeps based on H12 statistic present on islands but rare or absent on LVB mainland.

| Chr. | Region Start | Region End | Island Sites with Putative Sweep | Mainland Sites with Putative Sweep | Outlier Island Localities | Outlier Mainland Localities | Ag1000G Populations with Putative Sweep |
| --- | --- | --- | --- | --- | --- | --- | --- |
| 2R | 16,200,000 | 16,300,000 | 4 / 5 | 0 / 4 | Banda; Bukasa; Nsadzi; Sserinya | None | Guinea-Bissau |
| 2R | 17,300,000 | 17,500,000 | 4 / 5 | 1 / 4 | Banda; Bugala (I); Bukasa; Sserinya | Buwama | Guinea-Bissau |
| 2R | 21,000,000 | 21,100,000 | 5 / 5 | 1 / 4 | Banda; Bugala (I); Bukasa; Nsadzi; Sserinya | Buwama | None |
| 2R | 40,400,000 | 40,800,000 | 4 / 5 | 1 / 4 | Bugala (I); Bukasa; Nsadzi; Sserinya | Wamala | Burkina Faso [*gambiae*],<br>Cameroon [*gambiae*],<br>Gabon [*gambiae*], Guinea-Bissau |
| 2R | 41,100,000 | 41,200,000 | 4 / 5 | 1 / 4 | Banda; Bukasa; Nsadzi; Sserinya | Wamala | Cameroon [*gambiae*] |
| 2R | 55,800,000 | 55,900,000 | 4 / 5 | 1 / 4 | Banda; Bugala (I); Bukasa; Sserinya | Kiyindi | Angola [*coluzzii*] |
| 2L | 7,700,000 | 7,800,000 | 4 / 5 | 1 / 4 | Banda; Bugala (I); Bukasa; Nsadzi | Buwama | Guinea-Bissau, Uganda [*gambiae*] |
| 2L | 8,100,000 | 8,200,000 | 4 / 5 | 0 / 4 | Banda; Bugala (I); Nsadzi; Sserinya | None | None |
| 2L | 42,400,000 | 42,500,000 | 4 / 5 | 0 / 4 | Banda; Bugala (I); Nsadzi; Sserinya | None | None |
| 2L | 43,500,000 | 43,600,000 | 5 / 5 | 1 / 4 | Banda; Bugala (I); Bukasa; Nsadzi; Sserinya | Buwama | None |
| 2L | 49,000,000 | 49,100,000 | 4 / 5 | 1 / 4 | Banda; Bugala (I); Bukasa; Sserinya | Wamala | None |
| 3R | 26,600,000 | 26,700,000 | 5 / 5 | 0 / 4 | Banda; Bugala (I); Bukasa;<br>Nsadzi; Sserinya | None | None |
| 3R | 36,700,000 | 36,800,000 | 4 / 5 | 0 / 4 | Banda; Bugala (I); Bukasa;<br>Nsadzi | None | None |
| 3R | 44,200,000 | 44,300,000 | 4 / 5 | 1 / 4 | Banda; Bukasa; Nsadzi;<br>Sserinya | Kiyindi | Angola [*coluzzii*] |
| 3R | 46,200,000 | 46,300,000 | 4 / 5 | 0 / 4 | Banda; Bukasa; Nsadzi;<br>Sserinya | None | None |
| X | 6,600,000 | 7,000,000 | 4 / 5 | 0 / 4 | Banda; Bugala (I); Bukasa;<br>Nsadzi; Sserinya | None | Burkina Faso [*coluzzii*],<br>Gabon [*gambiae*], Guinea-Bissau |
| X | 8,100,000 | 10,700,000 | 4 / 5 | 0 / 4 | Banda; Bugala (I); Bukasa;<br>Nsadzi; Sserinya | Kiyindi | Burkina Faso [*coluzzii*],<br>Burkina Faso [*gambiae*],<br>Gabon [*gambiae*] |
| X | 11,300,000 | 11,800,000 | 5 / 5 | 0 / 4 | Banda; Bugala (I); Bukasa;<br>Nsadzi; Sserinya | None | Gabon [*gambiae*] |
| X | 12,900,000 | 13,000,000 | 4 / 5 | 0 / 4 | Banda; Bugala (I); Bukasa;<br>Sserinya | None | Gabon [*gambiae*] |
| X | 14,300,000 | 14,400,000 | 5 / 5 | 1 / 4 | Banda; Bugala (I); Bukasa;<br>Nsadzi; Sserinya | Kaazi | Gabon [*gambiae*] |
| X | 16,200,000 | 16,300,000 | 4 / 5 | 1 / 4 | Banda; Bugala (I); Bukasa;<br>Sserinya | Kaazi | Burkina Faso [*coluzzii*],<br>Gabon [*gambiae*] |

**Table S8:** Putative sweeps based on H12 statistic present on LVB mainland but rare or absent on islands.

| Chr. | Region Start | Region End | Island Sites with Putative Sweep | Mainland Sites with Putative Sweep | Outlier Island Localities | Outlier Mainland Localities | Ag1000G Populations with Putative Sweep |
| --- | --- | --- | --- | --- | --- | --- | --- |
| 2R | 27,600,000 | 27,700,000 | 1 / 5 | 3 / 4 | Nsadzi | Buwama; Kiyindi; Wamala | None |
| 2R | 38,000,000 | 38,100,000 | 1 / 5 | 3 / 4 | Bugala (I) | Buwama; Kiyindi; Wamala | None |
| 2R | 42,700,000 | 42,800,000 | 0 / 5 | 3 / 4 | None | Buwama; Kiyindi; Wamala | None |
| 2R | 45,400,000 | 45,500,000 | 1 / 5 | 3 / 4 | Sserinya | Buwama; Kiyindi; Wamala | None |
| 2R | 46,800,000 | 46,900,000 | 1 / 5 | 3 / 4 | Banda | Buwama; Kiyindi; Wamala | Cameroon [*gambiae*] |
| 2R | 48,000,000 | 48,100,000 | 1 / 5 | 3 / 4 | Bukasa | Buwama; Kaazi; Wamala | Angola [*coluzzii*],<br>Cameroon [*gambiae*] |
| 2R | 48,800,000 | 48,900,000 | 1 / 5 | 3 / 4 | Nsadzi | Buwama; Kaazi; Wamala | None |
| 2R | 50,900,000 | 51,000,000 | 1 / 5 | 3 / 4 | Bukasa | Kaazi; Kiyindi; Wamala | Burkina Faso [*gambiae*],<br>Guinea [*gambiae*] |
| 2R | 51,500,000 | 51,600,000 | 0 / 5 | 3 / 4 | None | Kaazi; Kiyindi; Wamala | None |
| 2R | 57,500,000 | 57,600,000 | 1 / 5 | 3 / 4 | Banda | Buwama; Kaazi; Kiyindi | Angola [*coluzzii*], Guinea-Bissau |
| 2L | 2,900,000 | 3,000,000 | 1 / 5 | 4 / 4 | Sserinya | Buwama; Kaazi; Kiyindi; Wamala | Angola [*coluzzii*], Burkina Faso [*coluzzii*], Burkina Faso [*gambiae*], Cameroon [*gambiae*], Gabon [*gambiae*], Guinea [*gambiae*], Uganda [*gambiae*] |
| 2L | 4,200,000 | 4,300,000 | 1 / 5 | 4 / 4 | Bugala (I) | Buwama; Kaazi; Kiyindi; Wamala | Angola [*coluzzii*], Cameroon [*gambiae*], Gabon [*gambiae*], Uganda [*gambiae*] |
| 2L | 5,700,000 | 5,800,000 | 1 / 5 | 3 / 4 | Bugala (I) | Buwama; Kaazi; Kiyindi | Angola [*coluzzii*], Guinea [*gambiae*], Uganda [*gambiae*] |
| 2L | 6,200,000 | 6,300,000 | 1 / 5 | 3 / 4 | Bugala (I) | Kaazi; Kiyindi; Wamala | Uganda [*gambiae*] |
| 2L | 6,600,000 | 6,800,000 | 1 / 5 | 3 / 4 | Bugala (I) | Kaazi; Kiyindi; Wamala | Angola [*coluzzii*], Cameroon [*gambiae*], Gabon [*gambiae*], Guinea [*gambiae*], Uganda [*gambiae*] |
| 2L | 10,000,000 | 10,100,000 | 1 / 5 | 3 / 4 | Sserinya | Kaazi; Kiyindi; Wamala | None |
| 2L | 10,800,000 | 10,900,000 | 0 / 5 | 3 / 4 | None | Kaazi; Kiyindi; Wamala | None |
| 2L | 11,300,000 | 11,400,000 | 1 / 5 | 3 / 4 | Bugala (I) | Kaazi; Kiyindi; Wamala | Guinea-Bissau |
| 2L | 12,000,000 | 12,100,000 | 1 / 5 | 3 / 4 | Bugala (I) | Kaazi; Kiyindi; Wamala | None |
| 2L | 12,400,000 | 13,000,000 | 0 / 5 | 3 / 4 | Bukasa | Buwama; Kaazi; Kiyindi; Wamala | None |
| 2L | 14,500,000 | 14,900,000 | 1 / 5 | 3 / 4 | Sserinya | Buwama; Kiyindi; Wamala | Gabon [*gambiae*], Uganda [*gambiae*] |
| 2L | 16,000,000 | 16,300,000 | 1 / 5 | 3 / 4 | Bukasa | Buwama; Kaazi; Wamala | Gabon [*gambiae*] |
| 2L | 16,600,000 | 16,700,000 | 1 / 5 | 4 / 4 | Bugala (I) | Buwama; Kaazi; Kiyindi; Wamala | None |
| 2L | 18,700,000 | 18,800,000 | 1 / 5 | 3 / 4 | Nsadzi | Kaazi; Kiyindi; Wamala | None |
| 2L | 33,600,000 | 33,700,000 | 1 / 5 | 3 / 4 | Bugala (I) | Buwama; Kaazi; Kiyindi | Angola [*coluzzii*] |
| 2L | 34,400,000 | 34,500,000 | 1 / 5 | 3 / 4 | Sserinya | Buwama; Kaazi; Wamala | None |
| 3R | 28,500,000 | 28,700,000 | 1 / 5 | 4 / 4 | Sserinya | Buwama; Kaazi; Kiyindi; Wamala | Burkina Faso [*coluzzii*], Burkina Faso [*gambiae*], Cameroon [*gambiae*], Gabon [*gambiae*], Guinea [*gambiae*], Uganda [*gambiae*] |
| 3R | 36,500,000 | 36,900,000 | 0 / 5 | 3 / 4 | Nsadzi | Buwama; Kiyindi; Wamala | None |
| 3R | 43,000,000 | 43,100,000 | 0 / 5 | 3 / 4 | None | Buwama; Kiyindi; Wamala | None |
| 3R | 43,700,000 | 44,100,000 | 0 / 5 | 3 / 4 | Nsadzi | Buwama; Kiyindi; Wamala | Angola [*coluzzii*], Burkina Faso [*gambiae*], Guinea [*gambiae*], Uganda [*gambiae*] |
| 3R | 48,800,000 | 48,900,000 | 0 / 5 | 3 / 4 | None | Buwama; Kiyindi; Wamala | None |
| 3R | 50,000,000 | 50,100,000 | 1 / 5 | 3 / 4 | Sserinya | Kaazi; Kiyindi; Wamala | None |
| 3L | 7,000,000 | 7,100,000 | 1 / 5 | 4 / 4 | Sserinya | Buwama; Kaazi; Kiyindi; Wamala | None |
| 3L | 11,500,000 | 11,600,000 | 1 / 5 | 3 / 4 | Sserinya | Buwama; Kiyindi; Wamala | Burkina Faso [*coluzzii*] |
| 3L | 12,200,000 | 12,300,000 | 0 / 5 | 3 / 4 | None | Kaazi; Kiyindi; Wamala | None |
| 3L | 13,400,000 | 13,500,000 | 0 / 5 | 3 / 4 | None | Kaazi; Kiyindi; Wamala | None |
| 3L | 16,300,000 | 16,400,000 | 1 / 5 | 3 / 4 | Sserinya | Buwama; Kiyindi; Wamala | Uganda [*gambiae*] |

**Table S9:** Signatures of selective sweeps on known insecticide genes by site based on H12 statistic.

| Chr. | Location | Insecticide<br>Gene | Island Sites with<br>Putative Sweep | Mainland Sites with<br>Putative Sweep | Outlier<br>Island Localities | Outlier<br>Mainland Localities | Ag1000G Populations<br>with Putative Sweep |
| --- | --- | --- | --- | --- | --- | --- | --- |
| 2R | 28,497,407 | Cyp6p | 5 / 5 | 4 / 4 | Banda; Bugala<br>(I); Bukasa;<br>Nsadzi; Sserinya | Buwama; Kaazi;<br>Kiyindi; Wamala | Angola [*coluzzii*], Burkina Faso<br>[*coluzzii*], Burkina Faso [*gambiae*],<br>Cameroon [*gambiae*], Guinea [*gam-*<br>*biae*], Uganda [*gambiae*] |
| 3R | 28,598,038 | Gste | 1 / 5 | 4 / 4 | Sserinya | Buwama; Kaazi;<br>Kiyindi; Wamala | Burkina Faso [*coluzzii*], Burkina<br>Faso [*gambiae*], Cameroon [*gam-*<br>*biae*], Gabon [*gambiae*], Guinea<br>[*gambiae*], Uganda [*gambiae*] |
| X | 15,241,718 | Cyp9k1 | 3 / 5 | 4 / 4 | Banda; Bukasa;<br>Sserinya | Buwama; Kaazi;<br>Kiyindi; Wamala | Burkina Faso [*coluzzii*], Burkina<br>Faso [*gambiae*], Gabon [*gambiae*],<br>Guinea [*gambiae*] |

**Table S10:** Software and versions used for major parts of analysis.

| Software | Version | Citation |
| --- | --- | --- |
| ea-utils | - | [64] |
| BWA | 0.7.16a | [67] |
| GATK | 3.8 | [68] |
| PLINK | 1.90b4.6 | [69, 70] |
| SHAPEIT2 | 2.837 | [71] |
| SAMtools/BCFtools | 1.5 | [72, 90] |
| ADMIXTURE | 1.3.0 | [20] |
| CLUMPAK | - | [73] |
| VCFtools | 0.1.15 | [86] |
| $\delta a\delta i$ (python package) | 1.7.0 | [82, 83] |
| Stairway plot - Jpopgen | 2-beta | [21] |
| selscan | 1.2.0a | [87] |
| adegenet (R package) | 2.1.0 | [91] |
| ape (R package) | 5.0 | [92] |
| RColorBrewer (R package) | 1.1-2 | [93] |
| dendextend (R package) | 1.6.0 | [94] |
| rehh (R package) | 2.0.2 | [95] |
| eigensoft | 7.2.1 | [75, 76] |
| GNU parallel | 20170422 | [96] |
| tabix | 1.5 | [90] |
| bedtools | 2.26.0 | [97] |

**Table S11:**
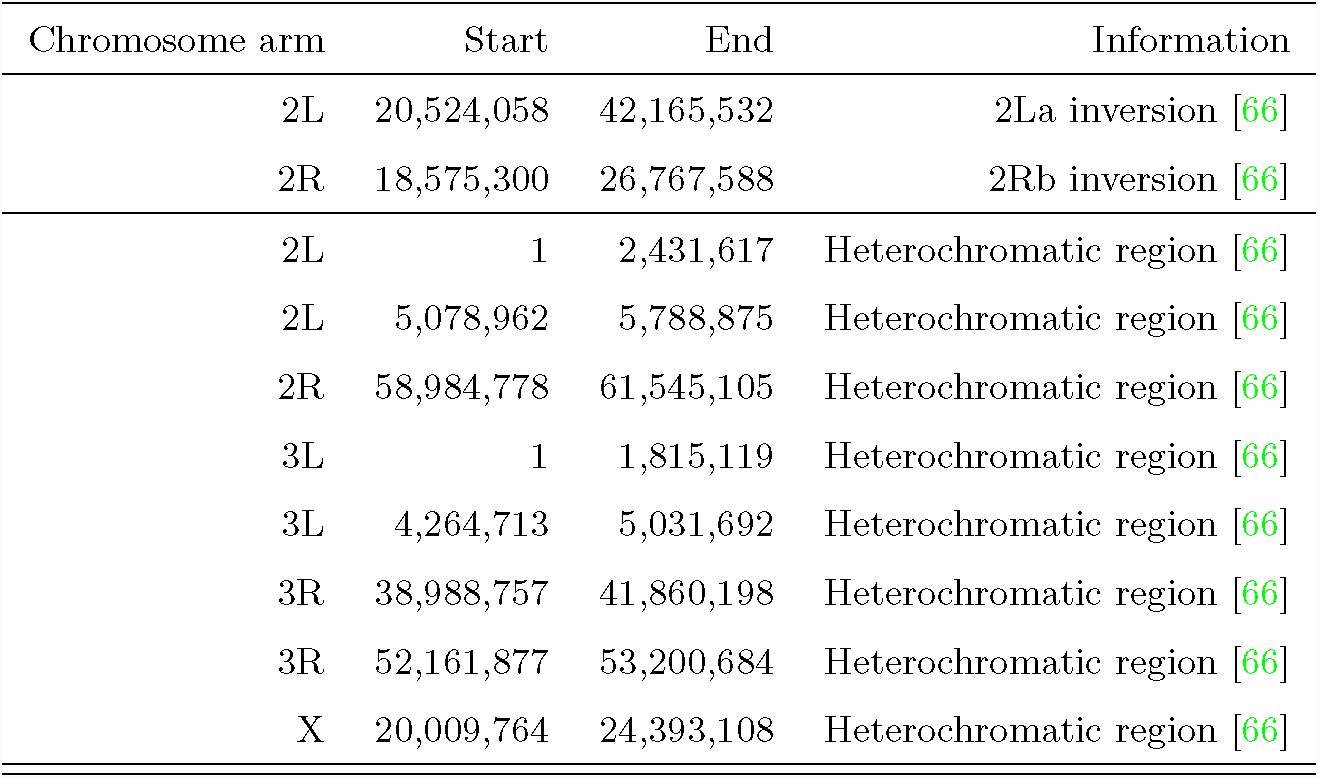
Genomic coordinates of heterochromatic and inverted regions.

| Chromosome arm | Start | End | Information |
| --- | --- | --- | --- |
| 2L | 20,524,058 | 42,165,532 | 2La inversion [66] |
| 2R | 18,575,300 | 26,767,588 | 2Rb inversion [66] |
| 2L | 1 | 2,431,617 | Heterochromatic region [66] |
| 2L | 5,078,962 | 5,788,875 | Heterochromatic region [66] |
| 2R | 58,984,778 | 61,545,105 | Heterochromatic region [66] |
| 3L | 1 | 1,815,119 | Heterochromatic region [66] |
| 3L | 4,264,713 | 5,031,692 | Heterochromatic region [66] |
| 3R | 38,988,757 | 41,860,198 | Heterochromatic region [66] |
| 3R | 52,161,877 | 53,200,684 | Heterochromatic region [66] |
| X | 20,009,764 | 24,393,108 | Heterochromatic region [66] |

